# Association between *ApoE* polymorphism and type 2 diabetes: A meta-analysis of 59 studies

**DOI:** 10.1101/530899

**Authors:** Dawei Chen, Jikang Shi, Yun Li, Yu Yang, Hui Yang, Shuping Ren

## Abstract

**Aims:** Due to the ever increasing incidence of T2DM, it is estimated that only half of the 79 million adults with type 2 diabetes (T2DM) will have adequate access to insulin by 2030 if the current levels of access is not improved. It is urgent to identify the important risk factors for T2DM and develop effective strategies to address the problem of T2DM. Our study aimed to evaluate the association between apolipoprotein E (*ApoE*) genetic polymorphism and type 2 diabetes, and to provide clues for the etiology of T2DM and even molecular marker of targeted therapy for the treatment of T2DM.

**Methods:** Case-control studies of ApoE polymorphism and T2DM, which were included in PubMed, Web of Science, Medline, WanFang, VIP, and CNKI databases, were selected and evaluated according to criteria of inclusion and exclusion. Eligible data were extracted and pooled, and were analyzed and assessed using R soft-ware (version 3.4.3). Random-effect models were used when heterogeneity existed in between-study, and fixed-effect models were applied otherwise.

**Results:** A total of 59 studies that consisted of 6,872 cases with T2DM and 8,250 controls were selected. Alleles and genotypes of *ApoE* between cases and controls were compared. For *ApoE* alleles, we observed the contrast of ε4 versus ε3 allele yielding a pooled OR of 1.18 (95% *CI:* 1.09-1.28; *P*<0.001). For *ApoE* genotypes, compared with ε3/ε3 genotype, ε2/ε2 genotype showed a possible association with T2DM (OR=1.46; 95% *CI:* 1.11-1.93; *P*=0.007), ε3/ε4 genotype had a 1.11-fold risk of developing T2DM (OR=1.11; 95% *CI:* 1.01-1.22; *P*=0.039), and ε4/ε4 genotype had a 1.71-fold risk of developing T2DM (*OR*=1.71; 95% *CI:* 1.33-2.19; *P*<0.001).

**Conclusions:** There is an association between *ApoE* polymorphism and T2DM: allele ε4 and genotypes (ε2/ε2, ε3/ε4, and ε4/ε4) are associated with the increased risk for the development of T2DM, and they may be risk factors for T2DM.

## 1. Introduction

It is estimated that only half of the 79 million adults with type 2 diabetes will have adequate access to insulin by 2030 if the current levels of access is not improved (Basu *et al.* 2019). Moreover, one of the significant causes of worldwide mortality and morbidity is diabetes (2016), especially type 2 diabetes mellitus, which is also the major cause of substantial global economic burden (Bommer *et al.* 2017). Therefore, there is an urgent need to identify the important risk factors for T2DM and develop effective strategies to address the problem of T2DM.

It is well accepted that genetic factor, environmental factors, and lifestyle contribute to the development of T2DM. Complex interactions between multiple genes and a range of environmental factors are involved in the onset and progression of type 2 diabetes(Scheuner *et al.* 2008). A better understanding of the contribution of genetic factors in the etiology of T2DM will facilitate the development of effective preventive strategies to reduce the ever increasing incidence of T2DM (Davies and Thirlaway 2013), it will also improve the effectiveness and precision of treatment and prevention strategies (O’Rahilly *et al.* 2005).

*ApoE* gene is one of the most studied genes which is responsible for stabilizing and solubilizing circulating lipoproteins in our body (Chaudhary *et al.* 2012). *ApoE* is a plasma glycoprotein of 34 kDa with 299-amino acids, and acts as a high affinity ligand for several hepatic lipoprotein receptors such as low-density lipoprotein receptor (LDLR) and LDL-related protein (LRP)(Chaudhary *et al.* 2012). *ApoE* is also involved in the process of cellular incorporation of several lipoproteins for transport and digestion (Mahley and Rall 2000) and is associated with several other plasma glycoproteins, such as high density lipoprotein (HDL), very low density lipoprotein (VLDL), and chylomicrons (Singh *et al.* 2006b). In humans, apoE gene is located on the chromosome at position 19q13.2 with SNPs at positions 112 (rs 429358) and 158 (rs 7412), and includes three major alleles: ε2 (T to C substitution at position 158), the most common ε3, and ε4 (C to T substitution at position 112); 3 isoforms: ApoE2 (Cys112, 158Cys), ApoE3 (Cys112, 158Arg), and ApoE4 (Arg112, 158Arg); and 6 genotypes having 3 homozygous: ε2/ε2, ε3/ε3, and ε4/ε4, and 3 heterozygous: ε2/ε3, ε2/ε4, and ε3/ε4 (Singh *et al.* 2006b).

*ApoE* is involved in many diseases, such as coronary heart disease (CHD))(Song *et al.* 2004), ischemic cerebrovascular disease (ICD)(Mccarron *et al.* 1999), Alzheimer’s disease (Farrer *et al.* 1997) and diabetes.

Much of the recent research has studied the association between the *ApoE* gene polymorphism and the risk of T2DM, however, there are inconsistencies between the results of the different studies. The inconsistency may result from the difference of included population, sample size, and genotyping methods. Moreover, 18 new papers(Chen 2006; Tang *et al.* 2007; Erdogan *et al.* 2009; Al-Majed *et al.* 2011; Chaudhary *et al.* 2012; Mustapic *et al.* 2012; Ge *et al.* 2013; Rong *et al.* 2013; Sun *et al.* 2013a; Xiong *et al.* 2013; Alharbi *et al.* 2014; Liu 2014; Wang *et al.* 2014; Atta *et al.* 2016; Liu *et al.* 2016; Luo *et al.* 2016; Mehmet *et al.* 2016; Liang *et al.* 2017) have been published since the publication of latest meta-analysis of the association between *ApoE* gene polymorphism and T2DM in 2014(Yin *et al.* 2014). Therefore, we enrolled these new published articles, and performed a further meta-analysis to investigate whether *ApoE* polymorphism is associated with the increased risk of T2DM.

## 2. Materials and Methods

### 2.1. Search strategy

We performed this meta-analysis by extensive literature search in PubMed, Web of Science, Medline, WanFang, VIP, and CNKI databases (last search on November 19, 2018). The terms used for searching were (“*ApoE*” OR “Apolipoprotein E”) AND (“polymorphism, Genetic” OR ‘‘variant” OR “mutation”) AND (“type 2 diabetes mellitus” OR “type 2 diabetes” OR “T2DM” OR “non-insulin dependent diabetes” OR “NIDDM”). The equivalent Chinese terms were used in the Chinese databases. In addition, we retrieved related articles that had not been identified in the initial search to replenish literatures.

### 2.2. Inclusion/exclusion criteria

Studies included in this meta-analysis were based on the following criteria: (1) case– control studies; (2) assessing the association between *ApoE* polymorphism and type 2 diabetes. The exclusion criteria met the follows: (1) duplicate articles; (2) no healthy controls; (3) lack of sufficient information on genotype or allele frequencies.

### 2.3. Data extraction

We extracted the main characteristics of each eligible study, including first author’s last name, date of publication, region, population’s ethnicity, genotyping method, number of cases and controls, and counts of the *ApoE* genotype or allele. We collected and calculated Hardy–Weinberg equilibrium (HWE) among the controls.

### 2.4. Quality assessment

The Newcastle-Ottawa scale (NOS) was used to evaluate quality of each article through a “star” rating system consisting of selection, comparability, and exposure. We allocated a score of 1 point for each condition a study met, and no point (0 score) if the condition or requirement was lacking. We calculated the total Quality Score of each study. Two authors (Jikang Shi and Shuping Ren) assessed the quality of included studies independently, When inconformity existed between the two authors, the results were requested to discuss with the third investigator (Dawei Chen). To avoid selection bias, studies with poor quality score were not excluded.

### 2.5. Statistical analysis

Allele and genotype frequencies of *ApoE* were calculated for each study to evaluate the HWE using Goodness of fit Chi-square test among control groups, and *P*<0.05 was considered as a significant deviation from HWE. The strength of association between *ApoE* polymorphisms and type 2 diabetes susceptibility was assessed using odds ratios (*OR*) and 95% confidence intervals (95% *CI*) because outcome variable was binary. Heterogeneity was evaluated by the Chi-square test based Q-statistic and quantified by *I^2^*-statistic (Higgins *et al.* 2003). Random-effect models (DerSimonian and Laird methods) were used to calculate *OR* and 95% *CI* when *P* value of *Q* test was more than 0.10 or *I*^2^ value was more than 50%; otherwise, fixed-effect models (Mantel and Haenszel methods) were applied (*I^2^*≥50% considered heterogeneity existed in between-study in this meta-analysis). Subgroup analyses stratified by ethnicity, quality score and Hardy–Weinberg equilibrium were performed to identify main sources of heterogeneity and to observe the association between *ApoE* polymorphisms and type 2 diabetes in different groups. Publication bias was assessed using funnel plots, and quantified by the Begg’s and Egger’s tests (*P*<0.05 considered statistically significant publication bias) (Begg and Mazumdar 1994). Sensitivity analysis was performed to examine stability of results by omitting each study in each turn. All data management and statistical analyses were used R soft-ware (version 3.4.3), *P*-value < 0.05 was considered statistically significant.

## 3. Results

### 3.1. Study Characteristics

Our meta-analysis initially collected 791 published articles, including 782 papers identified using our search strategy and 9 papers identified through the references. After abstracts and full texts were scanned according to the inclusion and exclusion criteria, 59 eligible articles with 6,872 cases and 8,250 controls were finally included in this paper. The protocol of the process for literature identification and selection is listed in Figure 1, and the baseline characteristics of the included studies are summarized in Table 1.

**Table 1.** Main characteristics of the included studies

| Study | Year | Region | Ethnicity | Genotyping method | Sample size<br>(case/control) | Quality score | HWE<br>Y/N(P) | $\epsilon 2/\epsilon 2(n)+\epsilon 2/\epsilon 3(n)$ | | $\epsilon 2/\epsilon 4(n)+\epsilon 3/\epsilon 3(n)$ | | $\epsilon 3/\epsilon 4(n)+\epsilon 4/\epsilon 4(n)$ | |
| --- | --- | --- | --- | --- | --- | --- | --- | --- | --- | --- | --- | --- | --- |
|  |  |  |  |  |  |  |  | case | control | case | control | case | control |
| Singh(SINGH *et al.* 2006a) | 2006 | India | Asian | PCR-RELP | 90/97 | 9 | Y(0.184) | 1+4 | 1+7 | 2+78 | 0+74 | 5+0 | 13+2 |
| Al-Majed(AL-MAJED *et al.* 2011) | 2011 | Kuwait | Other | PCR-RELP | 105/62 | 6 | N(0.006) | 7+2 | 2+3 | 2+73 | 2+46 | 6+15 | 9+1 |
| Chaudhary(CHAUDHARY *et al.* 2012) | 2012 | Bangkok | Other | PCR-RELP | 155/149 | 8 | Y(0.121) | 1+2 | 2+12 | 1+117 | 0+113 | 30+4 | 21+1 |
| Errera(ERRERA *et al.* 2006) | 2006 | Brazil | Other | PCR-RELP | 95/107 | 7 | Y(0.584) | 0+13 | 0+7 | 2+68 | 0+77 | 12+0 | 23+0 |
| Alharbi(ALHARBI *et al.* 2014) | 2014 | Riyadh | Other | TaqMan | 438/460 | 7 | N(<0.001) | 35+26 | 27+18 | 13+290 | 11+334 | 35+39 | 60+10 |
| Inamdar(INAMDAR *et al.* 2000) | 2000 | India | Asian | Flat gel isoelectric focusing | 60/40 | 8 | Y(0.054) | 2+8 | 1+9 | 3+17 | 2+10 | 16+14 | 8+10 |
| Kwon(KWON *et al.* 2007) | 2007 | Korea | Asian | PCR-RELP | 94/88 | 7 | Y(0.924) | 0+13 | 0+5 | 3+63 | 0+70 | 14+1 | 12+1 |
| Atta(ATTa *et al.* 2016) | 2016 | Egypt | Other | PCR-RELP | 45/45 | 5 | Y(0.098) | 0+12 | 0+3 | 12+12 | 3+30 | 9+0 | 9+0 |
| Vauhkonen(VAUHKONEN *et al.* 1997) | 1997 | Finland | Caucasian | PCR-RELP | 86/125 | 8 | Y(0.963) | 0+7 | 0+9 | 3+48 | 2+76 | 20+8 | 33+5 |
| Erdogan(ERDOGAN *et al.* 2009) | 2009 | Turkey | Caucasian | PCR-RELP | 56/35 | 7 | N(<0.001) | 0+4 | 0+0 | 0+40 | 0+28 | 12+0 | 7+0 |
| Eto(ETO *et al.* 1986) | 1986 | Japan | Asian | Flat gel isoelectric focusing | 105/111 | 8 | Y(0.339) | 0+9 | 1+10 | 0+73 | 1+80 | 21+2 | 16+3 |
| Guan(GUAN *et al.* 2009) | 2009 | Hong Kong | Asian | PCR-LDR | 213/111 | 7 | Y(0.499) | 8+32 | 1+32 | 7+141 | 1+88 | 24+1 | 9+1 |
| Leiva(LEIVA *et al.* 2005) | 2005 | Chile | Other | PCR-RELP | 193/139 | 7 | Y(0.293) | 0+12 | 0+10 | 4+133 | 3+87 | 43+1 | 39+0 |
| Liu(LIU *et al.* 2003) | 2003 | Shanghai | Asian | PCR-RELP | 80/81 | 7 | Y(0.217) | 0+11 | 0+4 | 1+56 | 2+64 | 12+0 | 11+0 |
| Mehmet(MEHMET *et al.* 2016) | 2015 | Turkey | Caucasian | PCR-RELP | 100/50 | 8 | N(0.039) | 0+6 | 0+22 | 0+81 | 0+19 | 13+0 | 9+0 |
| Xie(XIE *et al.* 2011) | 2011 | Hainan | Asian | PCR-RELP | 60/20 | 7 | Y(0.936) | 0+13 | 1+3 | 4+8 | 2+8 | 19+16 | 5+1 |
| Mustapic(MUSTAPIC *et al.* 2012) | 2012 | Croatia | Caucasian | TaqMan | 196/456 | 6 | Y(0.331) | 0+35 | 1+48 | 2+127 | 2+328 | 30+2 | 76+1 |
| Santos(SANTOS *et al.* 2002) | 2002 | Mexico | Other | PCR-RELP | 36/22 | 8 | Y(0.423) | 0+0 | 1+2 | 0+32 | 1+10 | 3+1 | 8+0 |
| Kamboh(KAMBOH *et al.* 1995) | 1995 | USA | Caucasian | IEF-immunoblotting and PCR | 116/659 | 6 | Y(0.992) | 0+23 | 6+88 | 5+62 | 19+382 | 26+0 | 150+14 |
| Ng(NG *et al.* 2006) | 2016 | Hong Kong | Asian | Other | 386/200 | 6 | Y(0.168) | 4+53 | 1+32 | 5+282 | 6+142 | 39+3 | 19+0 |
| Eto(ETO *et al.* 1995) | 1995 | Japan | Asian | Flat gel isoelectric focusing | 281/576 | 8 | Y(0.609) | 1+25 | 2+35 | 1+192 | 4+414 | 55+7 | 111+10 |
| Morbois<br>Trabut(MORBOIS-TRABUT *et al.* 2006) | 2006 | France | Caucasian | PCR-RELP | 210/481 | 7 | Y(0.773) | 2+31 | 5+71 | 1+143 | 14+294 | 33+0 | 87+10 |
| Powell(POWELL *et al.* 2003) | 2003 | UK | Caucasian | PCR-RELP | 187/102 | 7 | Y(0.094) | 3+22 | 2+7 | 3+89 | 1+57 | 27+3 | 21+0 |
| Guangda(GUANGDA *et al.* 1999) | 1999 | Wuhan | Asian | PCR-RELP | 89/72 | 7 | Y(0.122) | 1+13 | 1+7 | 1+66 | 2+53 | 7+1 | 7+2 |
| Zhang(ZHANG *et al.* 2000) | 2000 | Zhejiang | Asian | PCR-RELP | 63/71 | 8 | N(0.009) | 0+7 | 0+5 | 0+50 | 3+56 | 6+0 | 6+0 |
| Zhang(ZHANG *et al.* 2003) | 2003 | Sichuan | Asian | PCR-RELP | 74/191 | 8 | Y(0.878) | 0+5 | 1+23 | 1+55 | 1+134 | 12+1 | 31+1 |
| Sun(SUN *et al.* 2013b) | 2013 | Beijing | Asian | PCR-RELP | 243/78 | 7 | Y(0.414) | 6+36 | 2+12 | 0+180 | 1+55 | 21+0 | 6+1 |
| Hua(HUA *et al.* 2006) | 2006 | Jiangsu | Asian | PCR-RELP | 50/60 | 8 | Y(0.190) | 2+4 | 0+7 | 4+68 | 2+75 | 20+2 | 13+3 |
| Guo(GUO *et al.* 2003) | 2003 | Tianjin | Asian | PCR-RELP | 40/52 | 7 | Y(0.739) | 0+4 | 0+5 | 2+23 | 1+39 | 9+2 | 6+1 |
| Liang(LIANG *et al.* 2017) | 2017 | guangdong | Asian | PCR-RELP | 44/374 | 6 | Y(0.816) | 1+3 | 5+57 | 1+31 | 6+267 | 7+1 | 38+1 |
| Shen(SHEN *et al.* 2002a) | 2002 | Shanghai | Asian | PCR-RELP | 106/110 | 7 | Y(0.577) | 1+7 | 1+12 | 2+84 | 4+74 | 11+1 | 18+1 |
| Zheng(ZHENG *et al.* 1998) | 1998 | Shanghai | Asian | PCR-RELP | 112/60 | 8 | Y(0.801) | 2+16 | 1+8 | 1+81 | 0+45 | 11+1 | 6+0 |
| Hua(HUA *et al.* 2004) | 2004 | Suzhou | Asian | PCR-RELP | 38/60 | 7 | Y(0.434) | 1+7 | 0+4 | 2+24 | 1+45 | 4+0 | 8+2 |
| Liu(LIU 2014) | 2014 | Kunming | Asian | PCR-RELP | 215/298 | 7 | N(<0.001) | 10+0 | 2+0 | 0+174 | 0+272 | 31+0 | 23+1 |
| Xiang(GUANGDA *et al.* 1999) | 1995 | Kunming | Asian | PCR-RELP | 125/50 | 7 | Y(0.715) | 2+16 | 0+4 | 0+78 | 1+38 | 26+3 | 6+1 |
| Chen(CHEN 2006) | 2006 | Fujian | Asian | PCR-RELP | 97/105 | 7 | Y(0.906) | 2+15 | 1+18 | 1+70 | 2+72 | 8+1 | 10+1 |
| Xiang(GUANGDA XIANG 1999) | 1999 | Wuhan | Asian | PCR-ASO | 130/50 | 8 | Y(0.715) | 3+14 | 0+4 | 1+85 | 1+38 | 24+3 | 6+1 |
| Shen(SHEN *et al.* 2002b) | 2002 | Fujian | Asian | PCR-RELP | 35/50 | 6 | Y(0.112) | 3+11 | 0+6 | 2+4 | 4+31 | 14+0 | 9+0 |
| Xiong(XIONG *et al.* 2013) | 2013 | Hannan | Asian | PCR-RELP | 121/112 | 8 | Y(0.991) | 0+15 | 1+13 | 1+72 | 2+72 | 31+2 | 22+2 |
| Zhou(ZHOU *et al.* 2005) | 2005 | Heilongjiang | Asian | PCR-RELP | 67/68 | 7 | Y(0.263) | 0+13 | 2+9 | 1+47 | 0+46 | 6+0 | 11+0 |
| Xiang(GUANGDA XIANG 2005) | 2005 | Wuhan | Asian | PCR-ASO | 101/95 | 7 | Y(0.438) | 1+10 | 1+10 | 1+65 | 1+65 | 20+4 | 15+3 |
| Long(JIANQIU LONG 1999) | 1999 | Shanghai | Asian | PCR-RELP | 67/135 | 7 | Y(0.124) | 0+15 | 0+18 | 3+36 | 4+101 | 12+1 | 12+0 |
| Liang(SHU LIANG 2005) | 2005 | Jiangsu | Asian | PCR-RELP | 145/90 | 8 | Y(0.592) | 0+17 | 0+12 | 6+102 | 2+68 | 18+2 | 8+0 |
| Gu(LIQUN GU 2004) | 2004 | jiangsu | Asian | PCR-RELP | 63/90 | 8 | Y(0.592) | 0+9 | 0+12 | 3+43 | 2+68 | 7+1 | 8+0 |
| Yang(XIANGJIU YANG 1995) | 1995 | Hubei | Asian | PCR-RELP | 125/50 | 7 | N(0.028) | 2+16 | 1+3 | 0+78 | 1+38 | 26+3 | 5+2 |
| Rong(RONG *et al.* 2013) | 2013 | Guangdong | Asian | PCR-RELP | 18/29 | 7 | Y(0.953) | 0+4 | 0+8 | 0+18 | 0+29 | 2+0 | 1+0 |
| Liu(LIU *et al.* 2016) | 2016 | Yunnan | Asian | PCR-RELP | 300/300 | 8 | N(<0.001) | 14+0 | 2+0 | 0+243 | 0+274 | 43+0 | 23+1 |
| Tang(TANG *et al.* 2007) | 2007 | Zhenan | Asian | PCR-RELP | 41/60 | 6 | Y(0.80) | 0+1 | 0+3 | 2+28 | 1+43 | 10+0 | 13+0 |
| Qiu(QIU 2008) | 2008 | Zhejiang | Asian | PCR-RELP | 129/110 | 8 | Y(0.481) | 0+14 | 1+18 | 3+95 | 2+76 | 14+3 | 11+2 |
| Guo(JINJING GUO 2007) | 2007 | Gansu | Asian | ARMS-PCR | 40/40 | 6 | Y(0.618) | 0+1 | 1+4 | 3+29 | 1+27 | 7+1 | 7+0 |
| Xiong(YU XIONG 2008) | 2008 | Wuhan | Asian | MultiARMS PCR | 316/512 | 6 | Y(0.744) | 2+18 | 3+48 | 6+230 | 9+359 | 47+13 | 87+6 |
| Ge(GE *et al.* 2013) | 2013 | Inner Mongolia | Asian | PCR-RELP | 200/210 | 7 | Y(0.544) | 3+35 | 8+40 | 2+86 | 8+103 | 73+1 | 47+4 |
| Xiang(QIAN XIANG 2010) | 2010 | Yunnan | Asian | PCR-RELP | 41/102 | 7 | Y(0.473) | 0+5 | 0+13 | 1+28 | 0+70 | 7+0 | 19+0 |
| Luo(LUO *et al.* 2016) | 2016 | Guangdong | Asian | PCR-RELP | 35/50 | 6 | N(0.005) | 0+3 | 0+2 | 1+28 | 3+38 | 2+1 | 7+0 |
| Zhang(GUANGWU ZHANG 2007) | 2007 | Zhejiang | Asian | PCR-RELP | 38/49 | 6 | N(0.015) | 0+2 | 0+1 | 0+32 | 2+39 | 3+1 | 7+0 |
| Wang(WANG *et al.* 2014) | 2014 | Guangdong | Asian | PCR-RELP | 57/55 | 8 | N(0.027) | 0+4 | 2+7 | 2+33 | 4+28 | 13+5 | 8+6 |
| Zhang(LI ZHANG 1999) | 2002 | Anhui | Asian | PCR-RELP | 56/76 | 5 | Y(0.631) | 0+3 | 1+7 | 1+40 | 2+55 | 11+1 | 11+1 |
| Xiong(BIN XIONG 2005) | 2005 | Gansu | Asian | PCR-RELP | 32/30 | 7 | Y(0.608) | 1+5 | 0+4 | 1+22 | 1+23 | 2+1 | 2+0 |
| Dai(QINGFU DAI 2000) | 2000 | Fujian | Asian | PCR-RELP | 32/90 | 8 | Y(0.253) | 0+5 | 0+14 | 0+23 | 1+64 | 3+1 | 9+2 |

**Figure 1.**
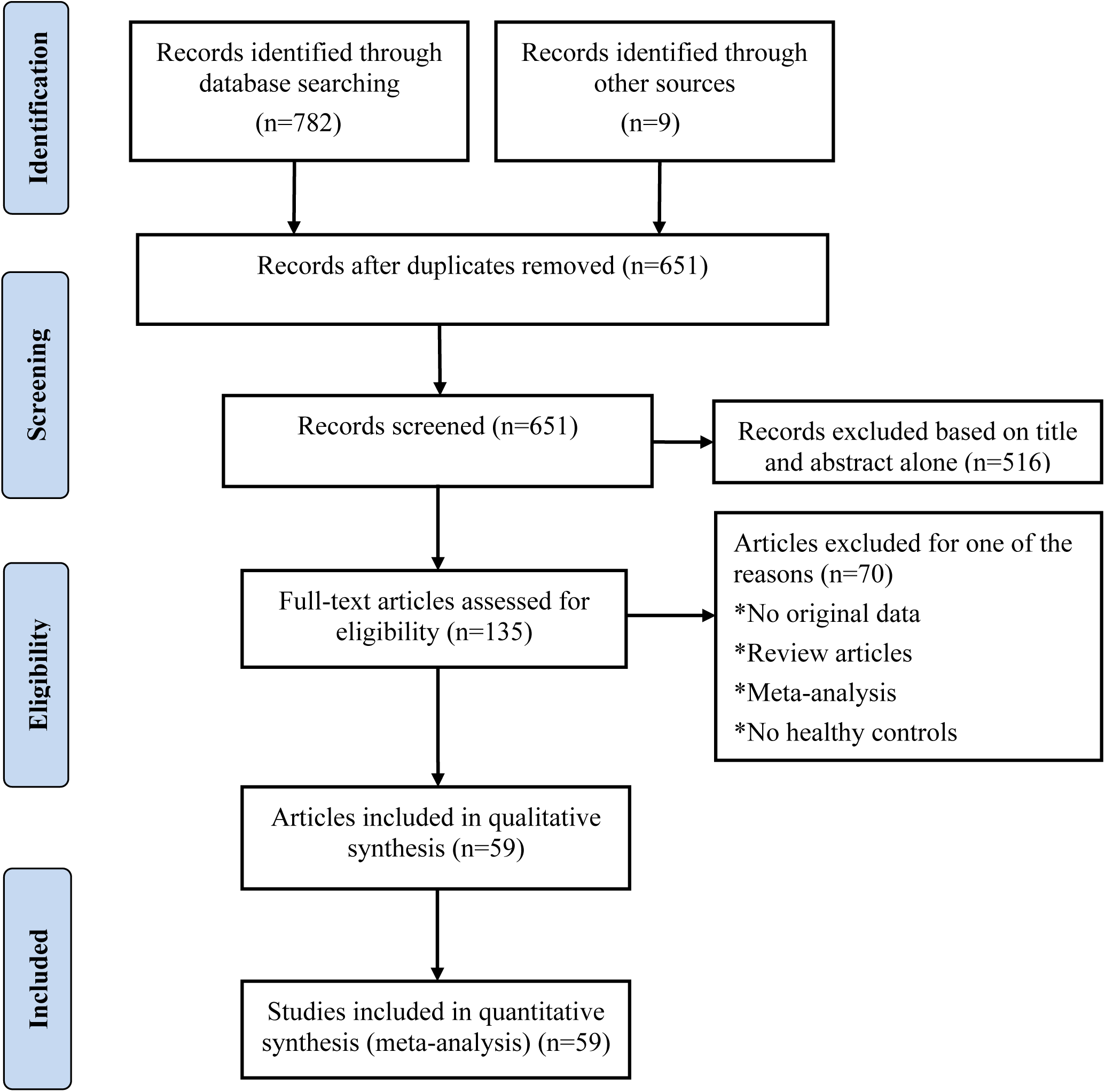
Flow chart of the process for literature identification and selection.

### 3.2. Association between alleles of *ApoE* and type 2 diabetes

There was significant heterogeneity in the comparison of *ApoE ε*2 with ε3 allele (*I*^2^=62%), and the pooled *OR* was 1.16 (95% *CI*: 0.98-1.37; *P*=0.079) when *ApoE ε*2 was compared with ε3 using the random-effects model (Figure 2); however, there was not heterogeneity in the comparison of *ApoE ε*4 with ε3 allele (*I*^2^=36%), and the pooled *OR* was 1.18 (95% *CI*: 1.09-1.28; *P*<0.001) when *ApoEε*4 was compared with ε3 using the fixed-effects model (Figure 3), suggesting that *ApoE ε*4 allele may be a risk factor for type 2 diabetes.

**Figure 2.**
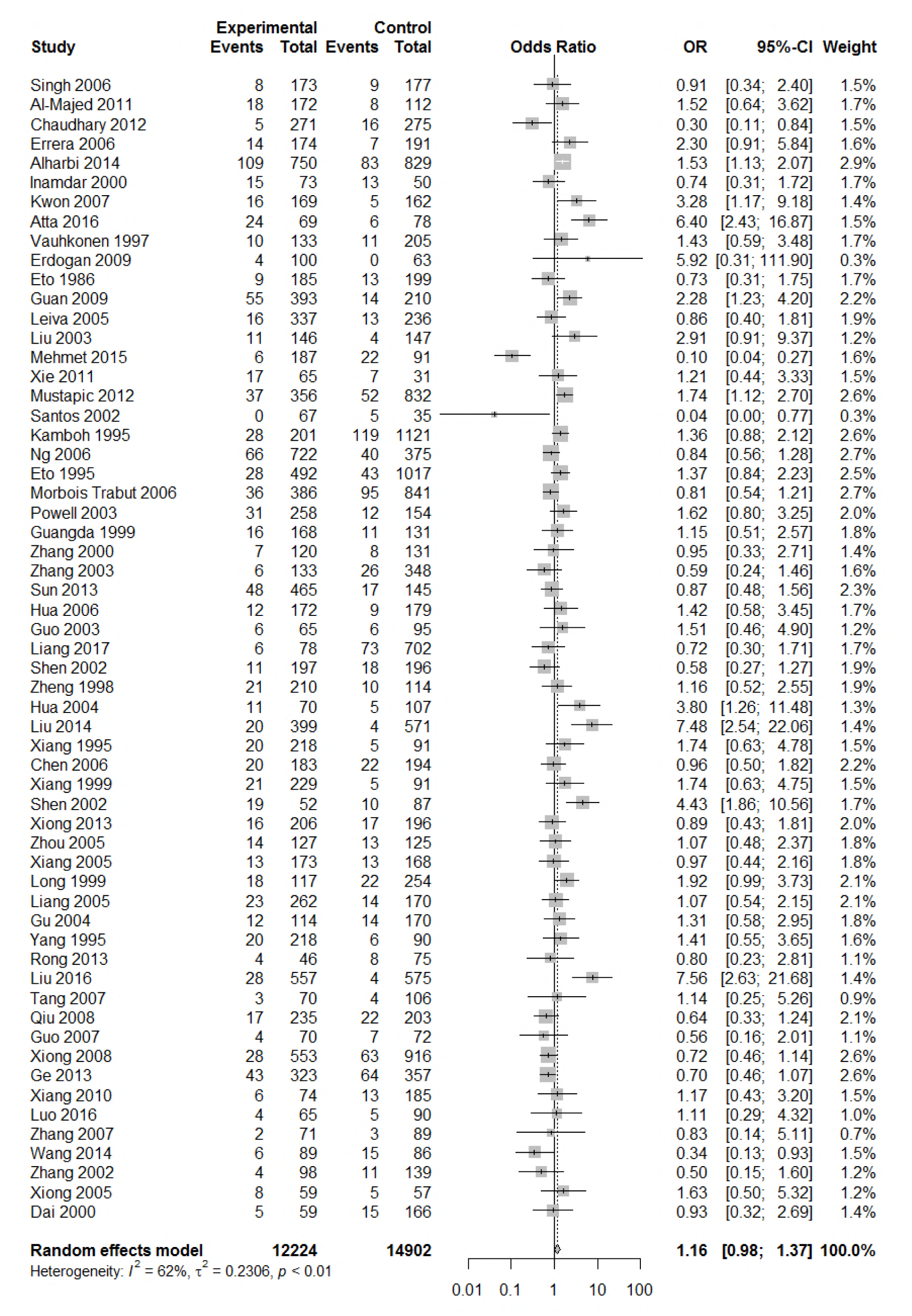
Forest plot for the result of association between type 2 diabetes and *ApoE ε*2 allele *vs*. ε3 allele based on a random-effects model.

**Figure 3.**
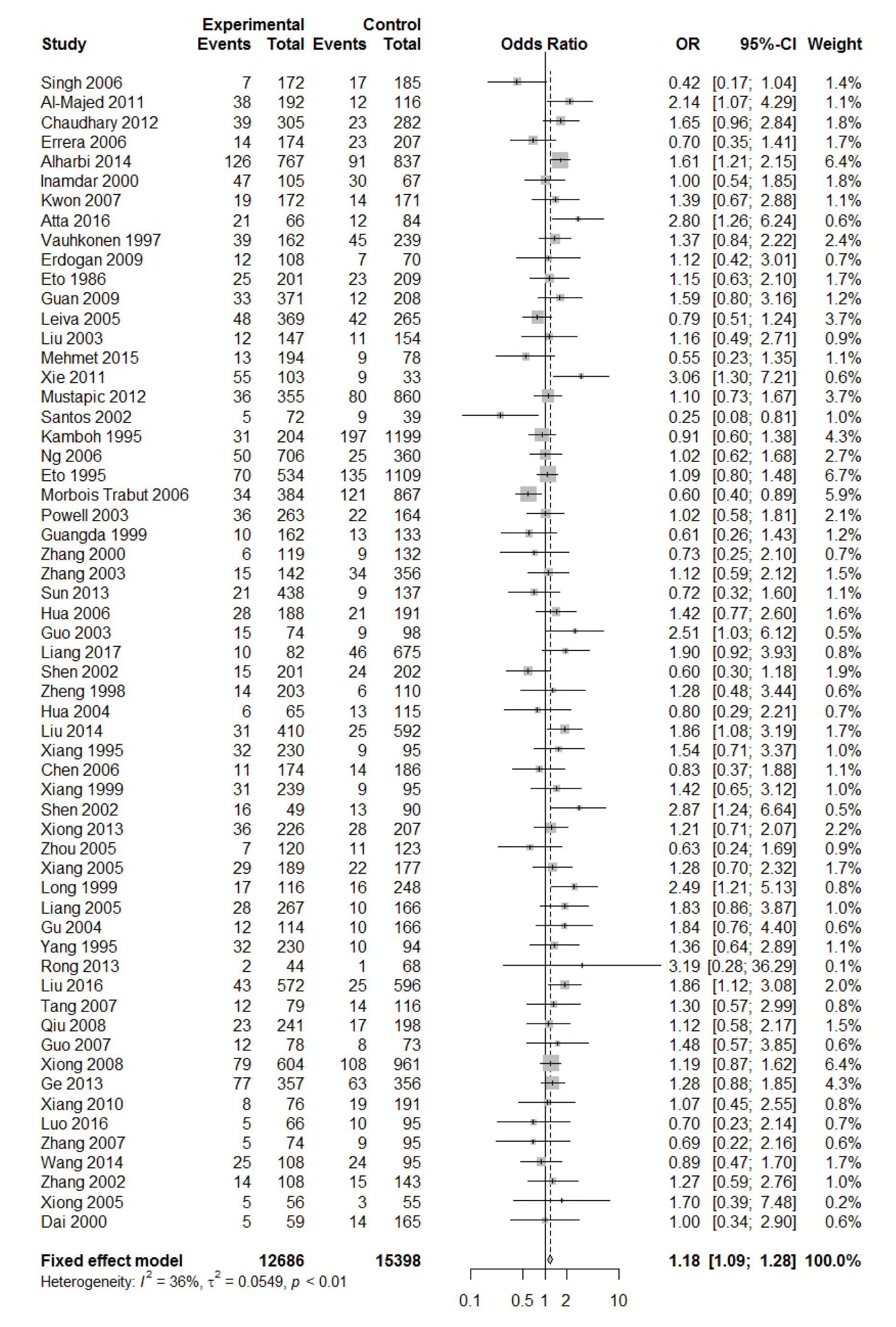
Forest plot for the result of association between type 2 diabetes and *ApoE ε*4 allele *vs*. ε3 allele based on a fixed-effects model.

### 3.3. Association between genotypes of *ApoE* and type 2 diabetes

There were five genotypes (ε2/ε2, ε2/ε3, ε2/ε4, ε3/ε*4*,and ε4/ε4) were compared with ε3/ε3 genotype. No significant heterogeneity was found whenthe ε2/ε2 genotype was compared with ε3/ε3 genotype (*I*^2^=0%), and the yielded *OR* of ε2/ε2 genotype versus ε3/ε3 genotype using a fixed-effects model was 1.46 (95% *CI:* 1.11-1.93; *P*=0.007) (Figure 4), suggesting that the ε2/ε2 genotype may have a harmful effect on type 2 diabetes. However, when ε2/ε3 genotype was compared with ε3/ε3 genotype, there was significant heterogeneity (*I*^2^=55%), and the yielded *OR* of ε2/ε3 genotype versus ε3/ε3 genotype using a random-effects model was 1.09 (95% *CI:* 0.90-1.32; *P*=0.397) (Figure 5). Compared with ε3/ε3 genotype, there were no significant heterogeneity between ε2/ε4, ε3/ε4, and ε4/ε4 genotype, respectively (*I*^2^=0%, *I*^2^=39%, and *I*^2^=0%). The yielded *OR* of ε2/ε4 genotype versus ε3/ε3 genotype using a fixed-effects model was 1.15 (95% *CI:* 0.90-1.46; *P*=0.276) (Figure 6). The yielded *OR* of ε3/ε4 genotype versus ε3/ε3 genotype using a fixed-effects model was 1.11 (95% *CI:* 1.01-1.22; *P*=0.039) (Figure 7). For the comparition of ε4/ε4 genotype with ε3/ε3 genotype, the yielded *OR* showed a 1.71-fold risk of type 2 diabetes (*OR*=1.7195% *CI:* 1.33-2.19; *P*<0.001) using the fixed-effects model (Figure 8).

**Figure 4.**
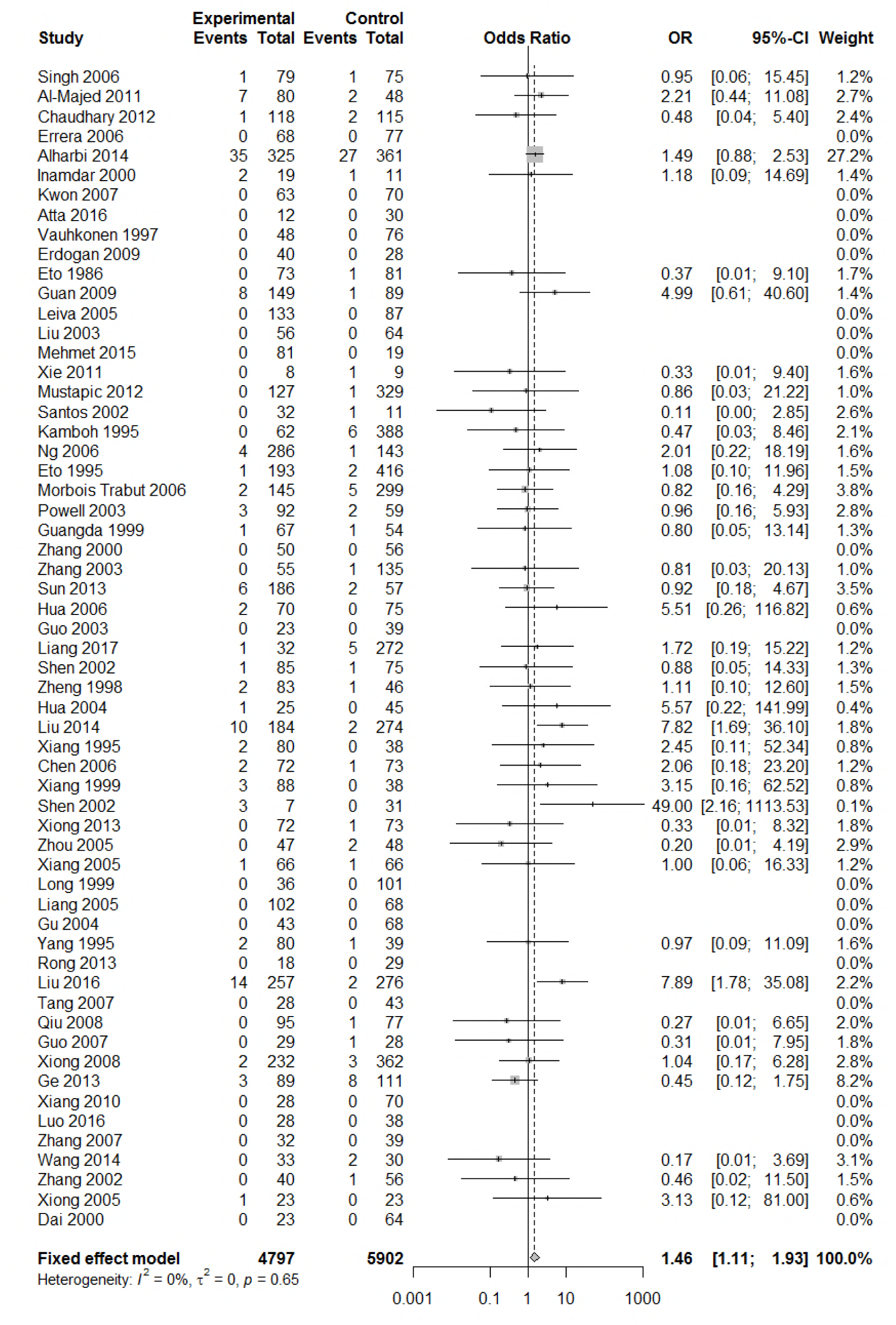
Forest plot for the result of association between type 2 diabetes and *ApoE ε*2/ε2 genotype *vs*. ε3/ε3 genotype based on a fixed-effects model.

**Figure 5.**
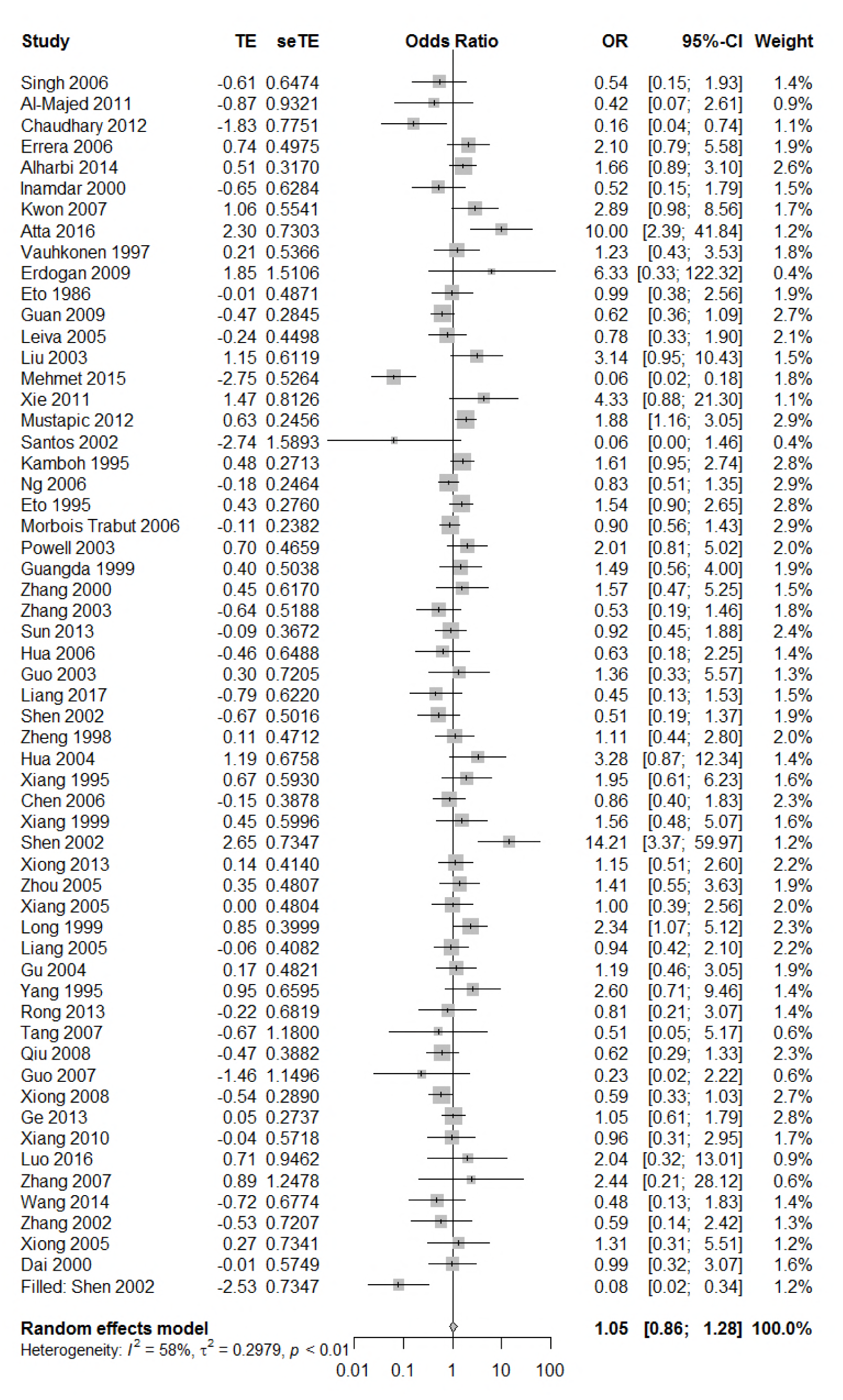
Forest plot for the result of association between type 2 diabetes and *ApoE ε*2/ε3 genotype *vs*. ε3/ε3 genotype based on a random-effects model.

**Figure 6.**
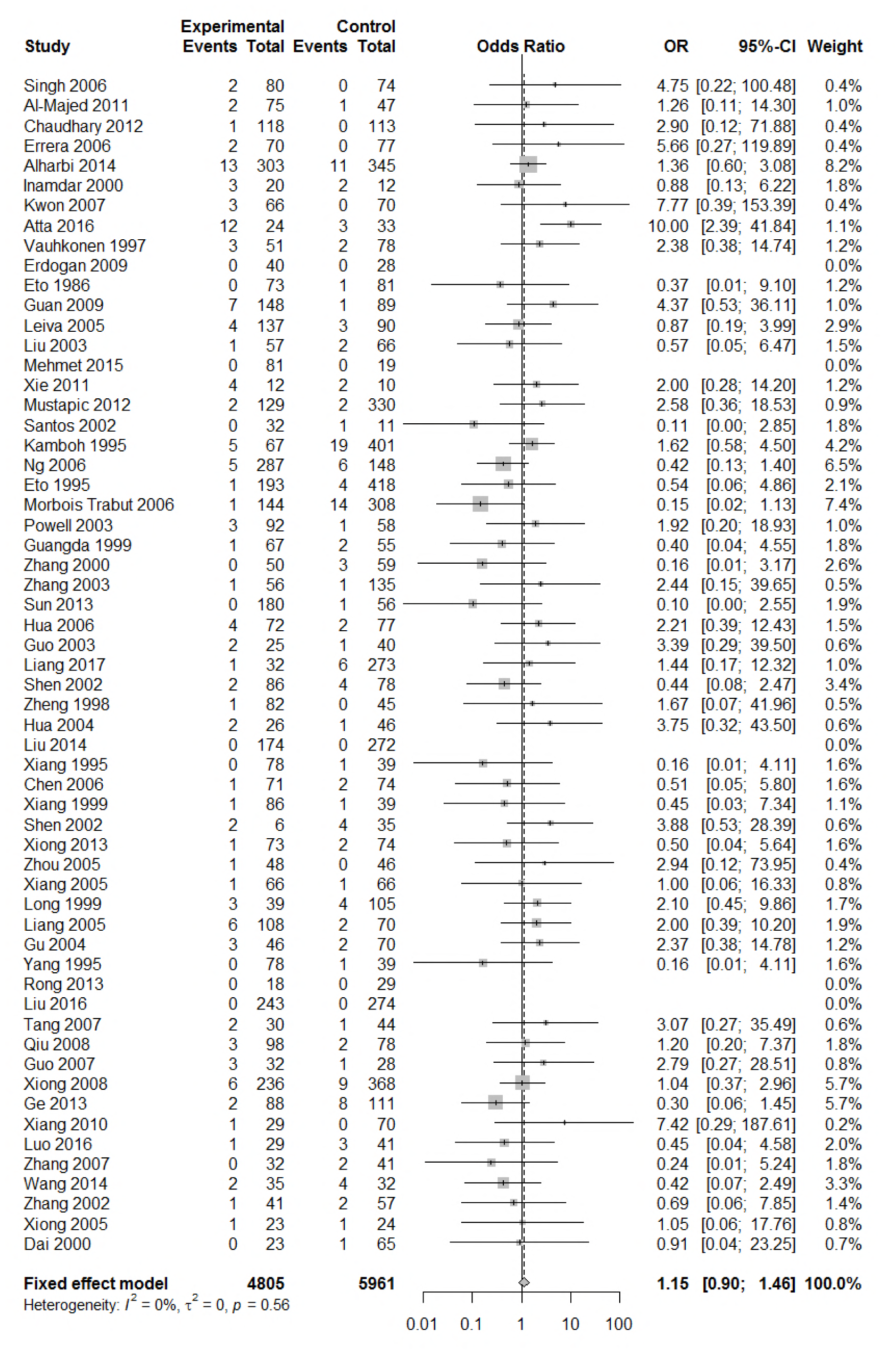
Forest plot for the result of association between type 2 diabetes and *ApoE ε*2/ε4 genotype *vs*. ε3/ε3 genotype based on a fixed-effects model.

**Figure 7.**
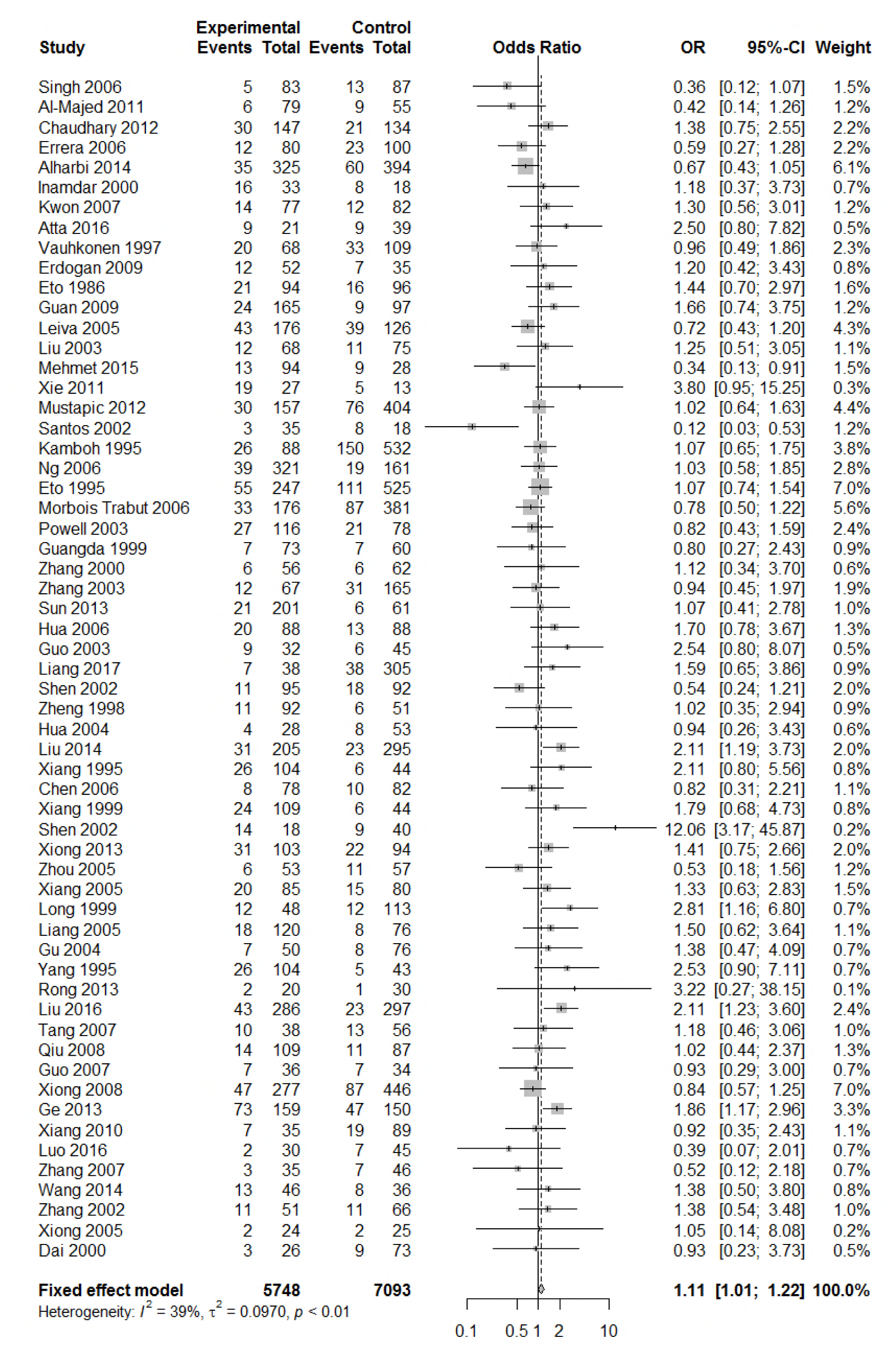
Forest plot for the result of association between type 2 diabetes and *ApoE ε*3/ε4 genotype *vs*. ε3/ε3 genotype based on a fixed-effects model.

**Figure 8.**
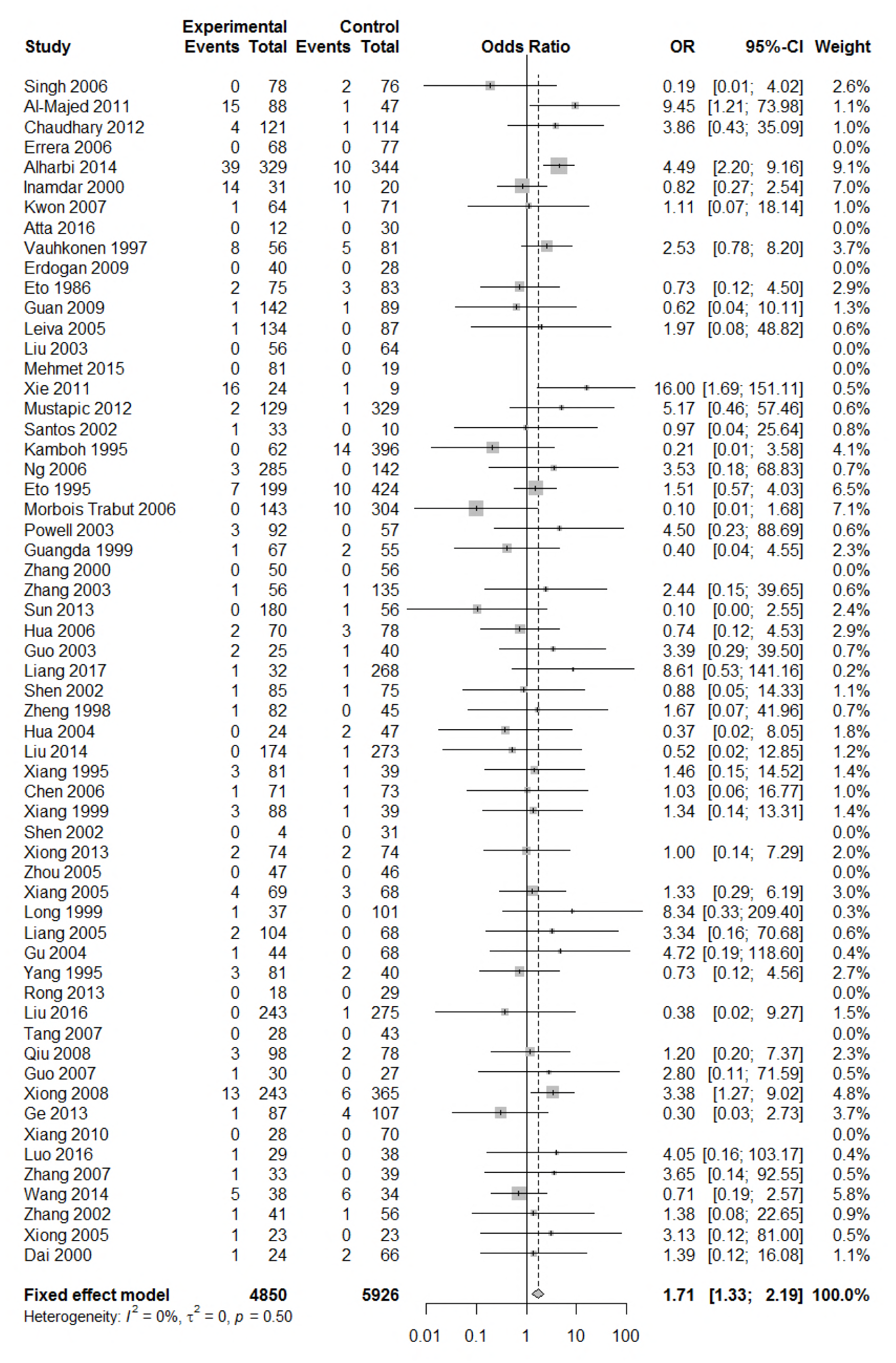
Forest plot for the result of association between type 2 diabetes and *ApoE ε*4/ε4 genotype *vs*. ε3/ε3 genotype based on a fixed-effects model.

### 3.4. Subgroup analysis

We conducted subgroup analysis stratified by ethnicity, quality score and Hardy– Weinberg equilibrium in order to identify main sources of heterogeneity. There were significant heterogeneity in the comparison of *ApoE ε*2 with ε3 allele (*I*^2^=62%) and the comparison of ε2/ε3 genotype with ε3/ε3 genotype (*I*^2^=55%) in our paper; however, we did not investigate sources of heterogeneity and there was no significant association between *ApoE* polymorphisms and type 2 diabetes in different subgroups (Supplementary Figure S1-S3).

### 3.5. Publication bias

Publication bias was assessed by funnel plots and quantified by Begg’s and Egger’s tests. All the funnel plots for *ApoE* allele and *ApoE* genotypes seemed symmetrical (Supplementary Figure S4-S5), and the results of Begg’s and Egger’s tests showed that there was no publication bias for the association between *ApoE* allele and type 2 diabetes and for the association between the *ApoE* genotypes and type 2 diabetes (all *P*>0.05).

### 3.6. Sensitivity analysis

Our results of sensitivity analysis showed that none of individual study influenced on the corresponding pooled *ORs* and 95% *CIs* in the comparison of *ApoE ε*4 with ε3 allele or in the comparison of *ApoE ε*2/ε3, ε2/ε4, and ε4/ε4 with genotype ε3/ε3 genotype (Figure 10, Figure 12, Figure 13, and Figure 15), suggesting that these results were relatively stable and credible. However, there were slight effects of individual study on the corresponding pooled *ORs* and 95% *CIs* in the comparison of *ApoE ε*2 with ε3 allele or in the comparison of *ApoE ε*2/ε2 and ε3/ε4 with genotype ε3/ε3 genotype (Figure 9, Figure 11, and Figure 14).

**Figure 9.**
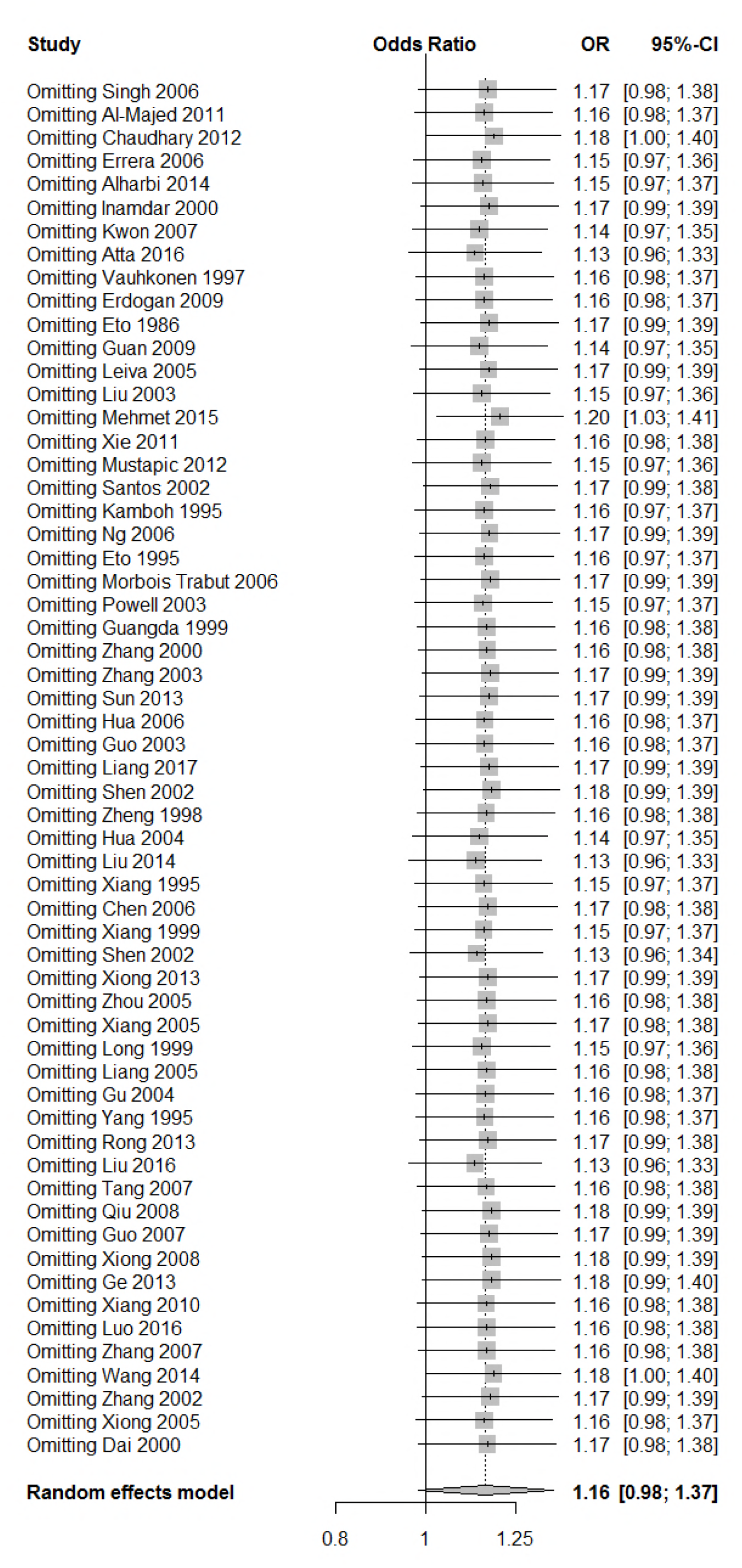
Sensitivity analysis for the result of association between type 2 diabetes and *ApoE ε*2 allele *vs*. ε3 allele.

**Figure 10.**
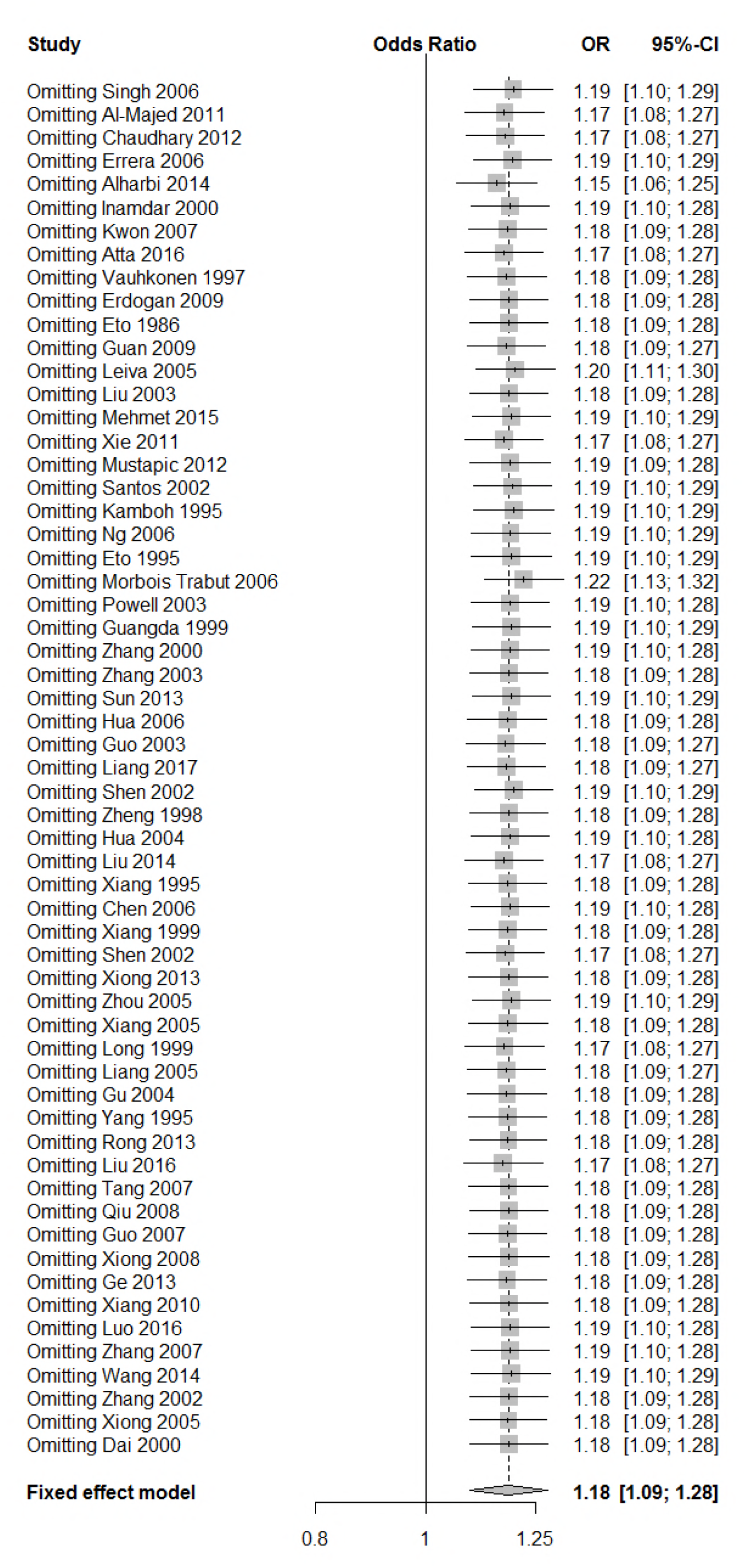
Sensitivity analysis for the result of association between type 2 diabetes and *ApoE ε*4 allele *vs*. ε3 allele.

**Figure 11.**
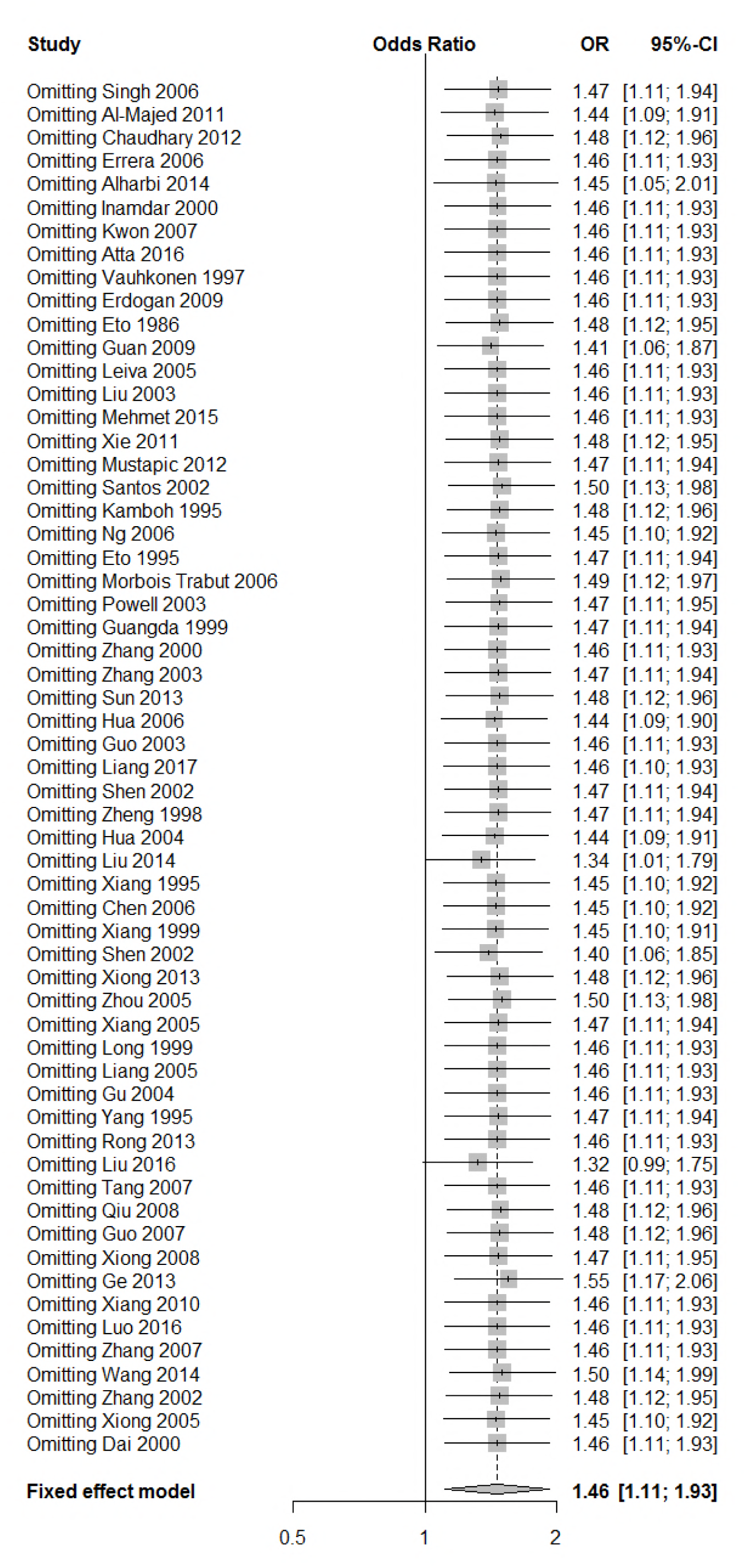
Sensitivity analysis for the result of association between type 2 diabetes and *ApoE ε*2/ε2 genotype *vs*. ε3/ε3 genotype.

**Figure 12.**
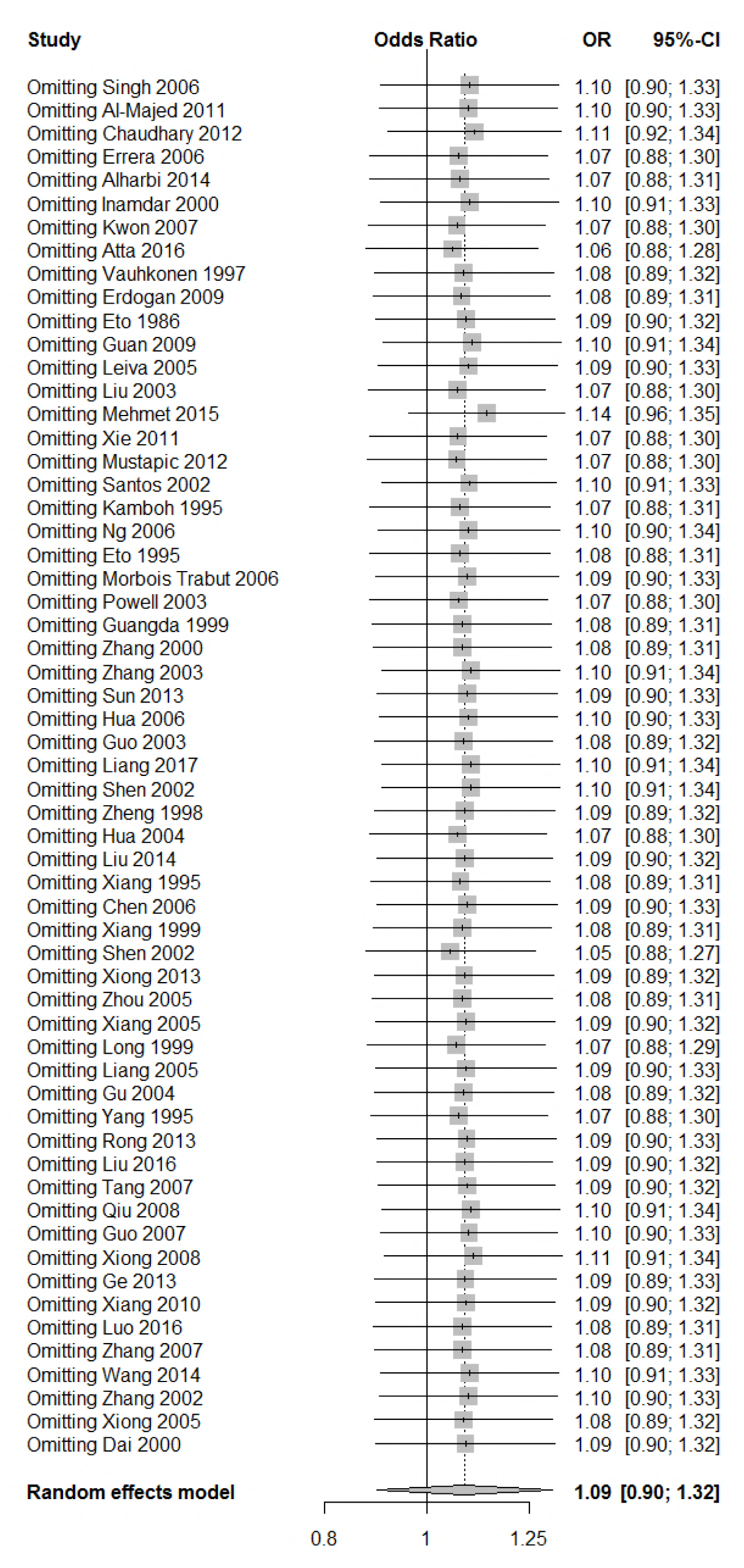
Sensitivity analysis for the result of association between type 2 diabetes and *ApoE ε*2/ε3 genotype *vs*. ε3/ε3 genotype.

**Figure 13.**
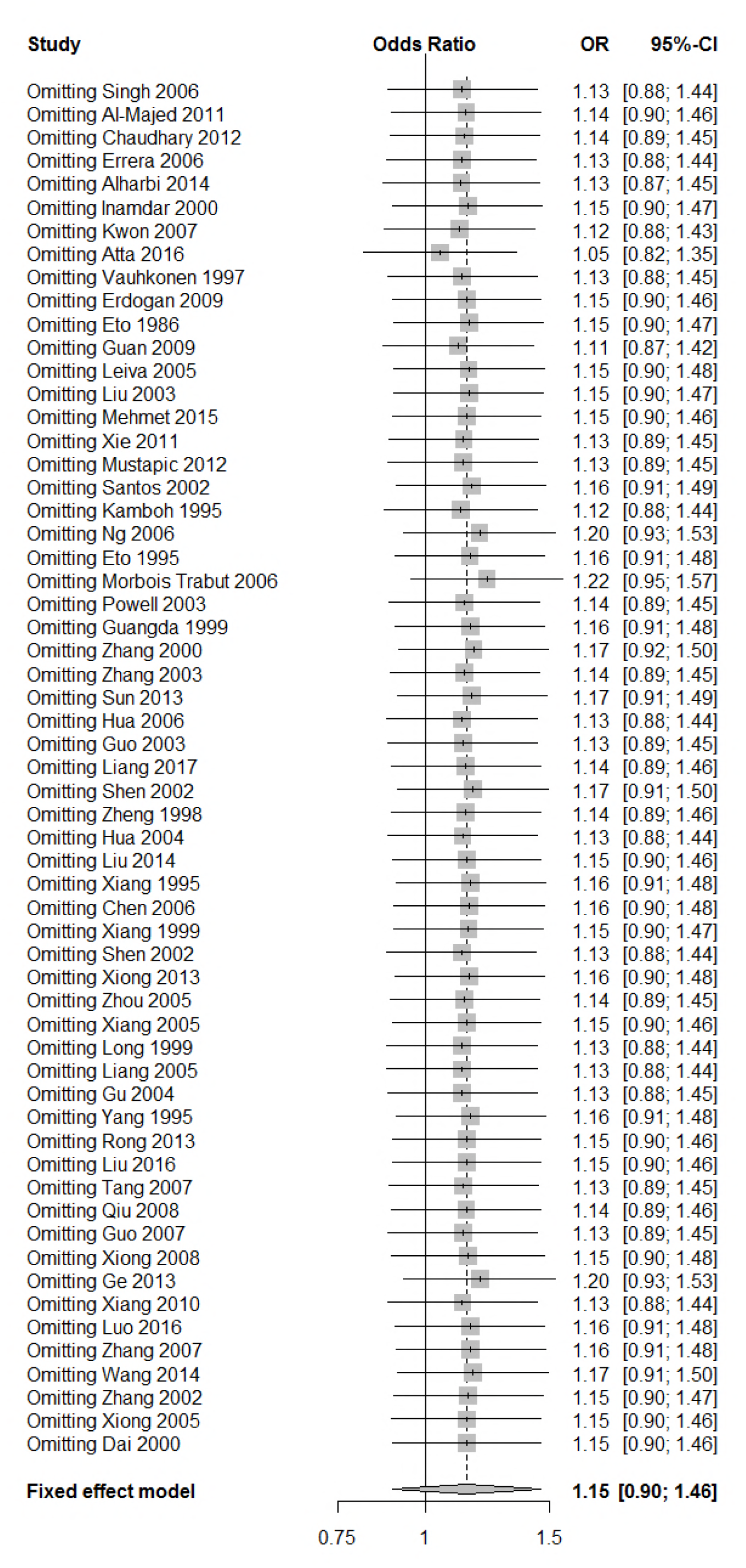
Sensitivity analysis for the result of association between type 2 diabetes and *ApoE ε*2/ε4 genotype *vs*. ε3/ε3 genotype.

**Figure 14.**
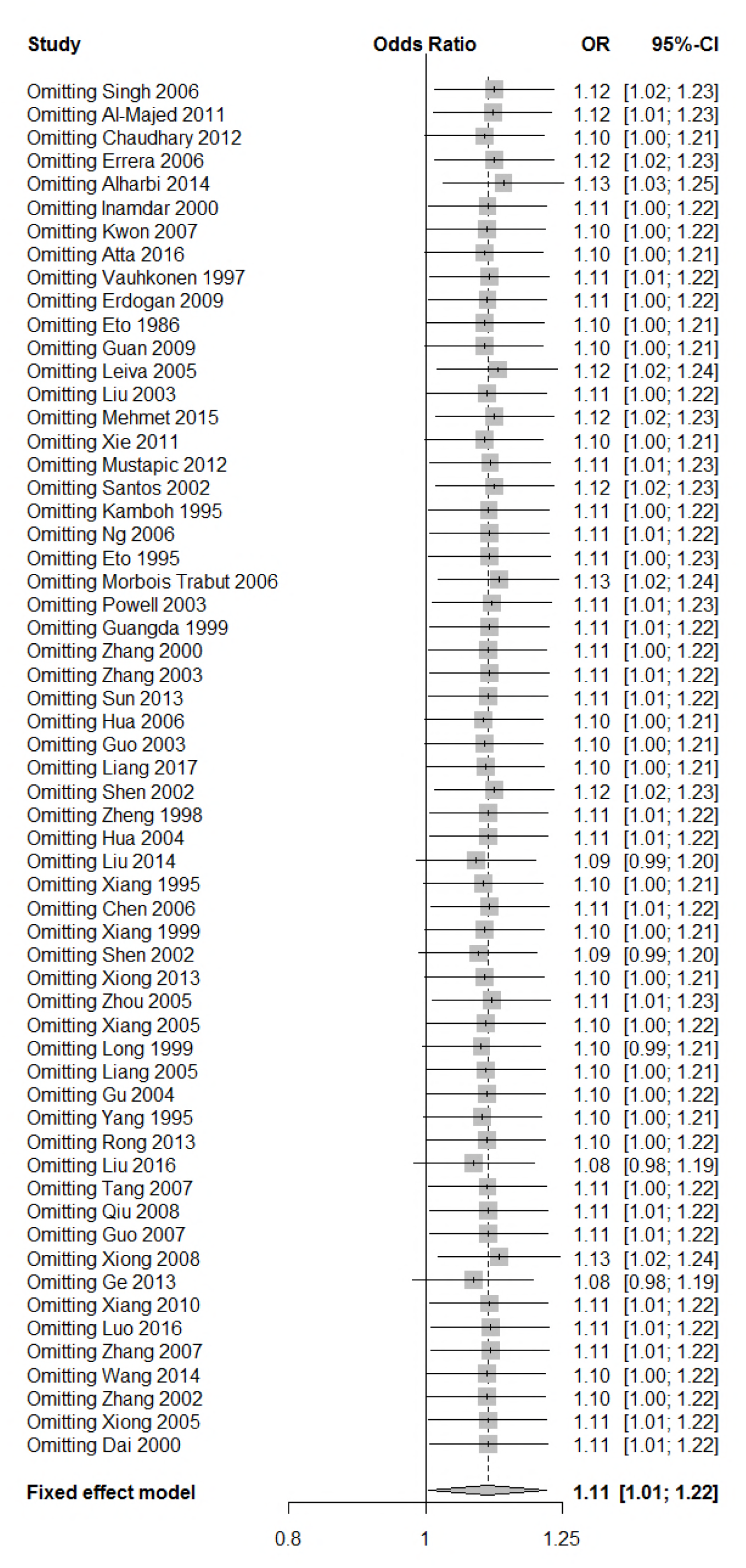
Sensitivity analysis for the result of association between type 2 diabetes and *ApoE ε*3/ε4 genotype *vs*. ε3/ε3 genotype.

**Figure 15.**
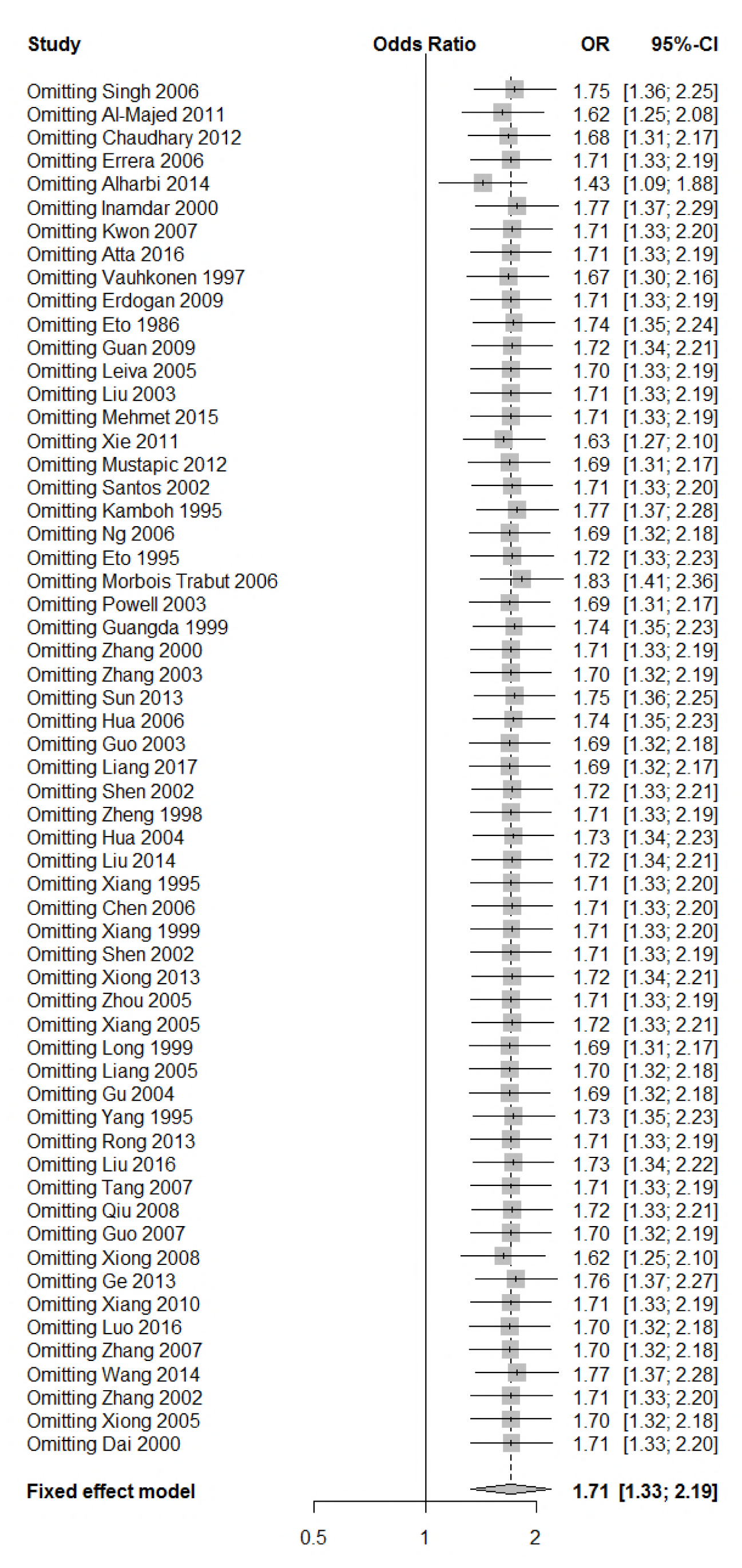
Sensitivity analysis for the result of association between type 2 diabetes and *ApoE ε*4/ε4 genotype *vs*. ε3/ε3 genotype.

## 4. Discussion

In this meta-analysis, we included 59 literatures with 6,872 cases and 8,250 controls to explore the association between the *ApoE* gene polymorphism and type 2 diabetes mellitus. The major findings of our study are that allele ε4 and genotypes (ε2/ε2, ε3/ε4, and ε4/ε4) are associated with the increased risk for the development of T2DM, however, allele ε2 and genotypes (ε2/ε3 and ε2/ε4) are not associated with T2DM.

The strengths of the present study are that, 1) we included all the published literatures on the association between *ApoE* gene polymorphism and T2DM regardless of regions or ethnicities; 2) we had a large sample size. There are 18 new published papers discussing the association between *ApoE* gene polymorphism and T2DM since the last meta-analysis published in 2014, all of them are included in our present meta-analysis, which will provide more convincing evidence to the association of *ApoE* gene polymorphism with T2DM; 3) the results of our sensitivity analysis demonstrate that the conclusion of the present study is very stable; 4) the results of publication bias analysis reveal that the conclusion of our study is absent of publication bias. However, our study also has several weaknesses, 1) presence of heterogenicity in our study. We did the subgroup analysis on HWE, genotyping methods and ethnicities, but we did not trace the source of heterogenicity; 2) since the present study is a case-control study, the findings of our study cannot provide the causal relationship between *ApoE* gene polymorphism and T2DM, only the association of *ApoE* gene polymorphism with T2DM.

The findings of our meta-analysis are in accordance with the previous studies(Anthopoulos *et al.* 2010; Qiu Xu 2010; Aimei Long 2013; Yin *et al.* 2014), showing that both *ApoE ε*4 allele and the genotypes (ε3/ε4 and ε4/ε4) were associated with increased risk of T2DM. Subjects carrying the ε4 alleles had higher plasma total cholesterol levels compared to subjects carrying the ε3/ε3 genotype, and HDL cholesterol was significantly lower in the ε3/ε4 than in the ε3/ε3 individuals(Dallongeville *et al.* 1992); individuals carrying the ε2/ε2 genotype had about 31% lower mean LDL than those with the ε4/ε4 genotype (Bennet *et al.* 2007). Insulin resistance is known to be strongly associated with metabolic dyslipidemia and the correlation of lipid profiles with diabetic phenotypes is significant. Therefore, *ApoE ε*4 allele and the genotypes (ε3/ε4 and ε4/ε4) were associated with an increased risk of T2DM through affecting the lipid metabolism.

We found the genotype ε2/ε2 was associated with increased risk of T2DM, but not allele ε2 or genotype ε2/ε3; which are not in agreement with the results of previous meta-analyses (Yin *et al.* 2014). The results from Yan et al’ showed that ε2 and genotype ε2/ε3 were associated with increased risk of T2DM, genotype ε2/ε2 was not associated with increased risk of T2DM. The inconsistency may be caused by the different subjects included. Yan et al’ research included only Chinese Han. Furthermore, we did not reveal the difference in the association of ApoE gene polymorphism with T2DM between ethnicities through subgroup analysis. In addition, our findings are consistent with those of Anthopoulos et al’ study (Anthopoulos *et al.* 2010) which reveals that the ORs for the other ε2-carriers genotypes (ε2/ε2, ε2/ε3, and ε2/ε4) compared to ε3/ε3 were greater than 1.00. The slight difference between the present study and Anthopoulos et al’ is that the OR of ε2/ε2 in our study reaches statistical significance while the OR of ε2/ε3 in Anthopoulos et al’ reaches statistical significance. However, the estimates of the results from Anthopoulos et al’ study are likely to be attenuated due to the small sample size. Our findings demonstrate that individuals with the genotype carrying single allele ε2 (ε2/ε3 and ε2/ε4) are not at the risk of T2DM while those carrying two ε2 allele (ε2/ε2) possess higher risk for T2DM, which also coincides with the finding that the higher frequency of the ε2/*APOE* allele might be primarily related to T2DM (Errera *et al.* 2006).

The significance of the present study is that we identified significant association between *ApoE* gene polymorphism and T2DM, which will provide clues for the etiology of T2DM and even molecular marker of targeted therapy for the treatment of T2DM. However, it is essential to further investigate the interaction between gene and gene as well as the gene and environment since T2DM is the result of interaction between genetic and environmental factors.

In conclusion, there is an association between *ApoE* polymorphism and T2DM: allele ε4 and genotypes (ε2/ε2, ε3/ε4, and ε4/ε4) are associated with the increased risk for the development of T2DM, and they may be risk factors for T2DM.

## Author Contributions

Conception and design: Shuping Ren. Provision of study materials: Dawei Chen, Jikang Shi, and Yun Li. Collection and assembly of data: Dawei Chen, Jikang Shi, Yun Li, and Yu Yang. Data analysis and interpretation: Jikang Shi and Hui Yang. Manuscript writing: Dawei Chen, Shuping Ren. Revised the language/article: All authors. Final approval of manuscript: All authors.

## Acknowledgements

This work was supported by the funds from Jipai Runda Environmental Inspection Technology Corporation Limited of Beijing (Grant No. 2015YX252) and Leshiguang Measurement Technology Corporation Limited (Grant No. 2018YX046).

---

**Supplementary Figure S1.**
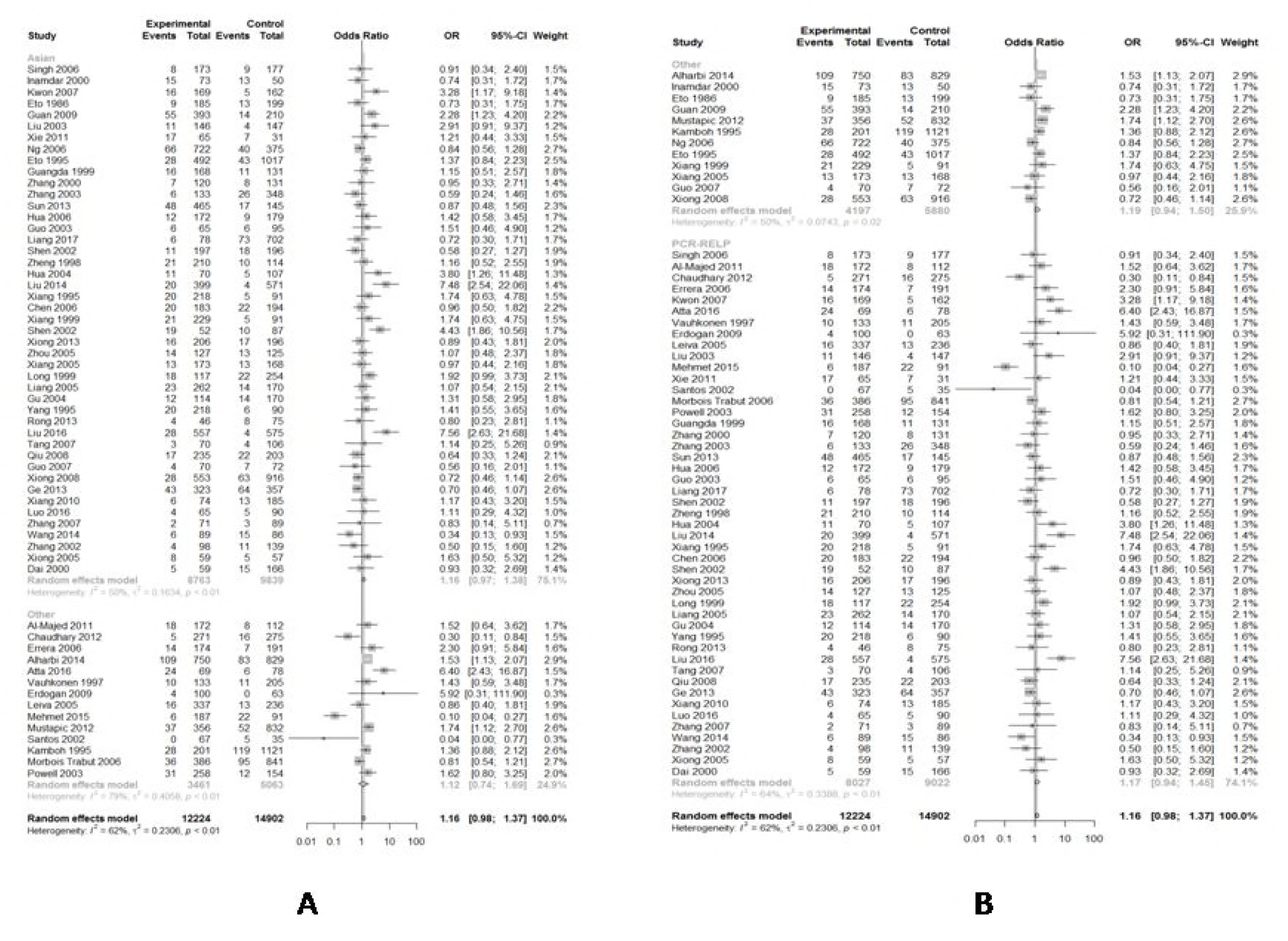
(A) Forest plot for associations between type 2 diabetes and *ApoE ε*2 allele *vs*. ε3 allele in the subgroup based on ethnicity. (B) Forest plot for associations between type 2 diabetes and *ApoE ε*2 allele *vs*. ε3 allele in the subgroup based on genotype.

**Supplementary Figure S2.**
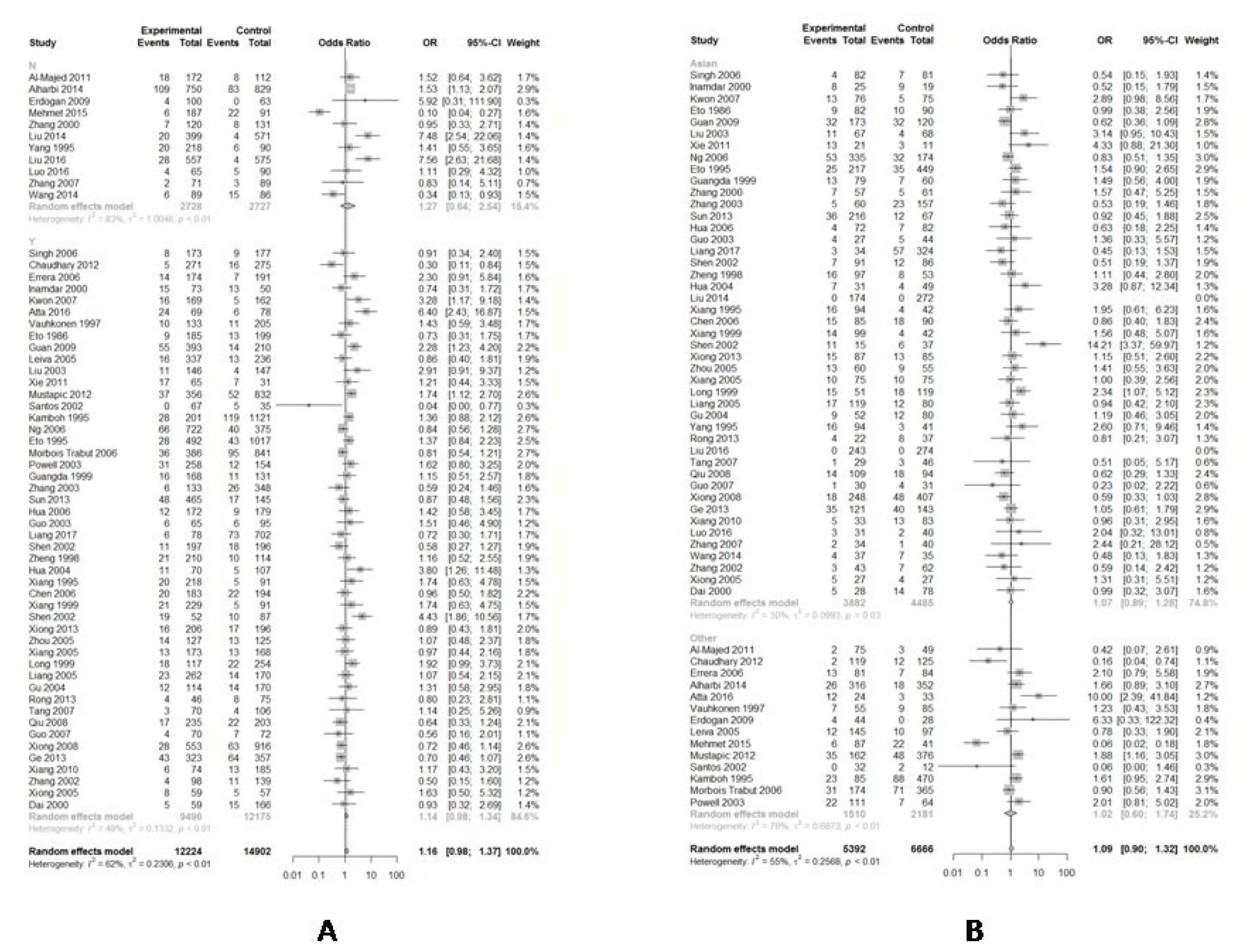
(A) Forest plot for associations between type 2 diabetes and *ApoE ε*2 allele *vs*. ε3 allele in the subgroup based on HWE. (B) Forest plot for associations between type 2 diabetes and *ApoE ε*2/ε3 genotype *vs*. ε3/ε3 genotype in the subgroup based on ethnicity.

**Supplementary Figure S3.**
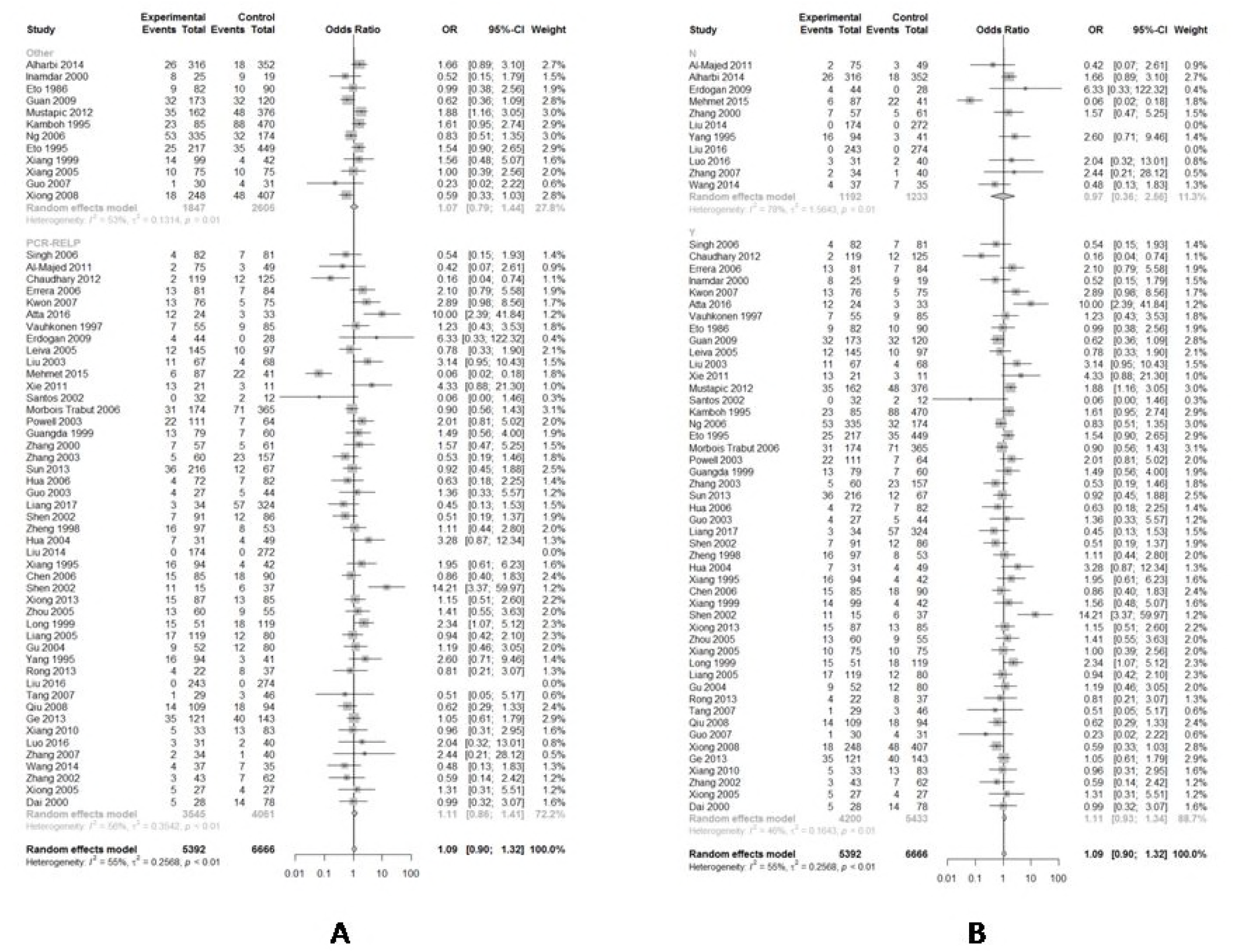
(A) Forest plot for associations between type 2 diabetes and *ApoE ε*2/ε3 genotype *vs*. ε3/ε3 genotype in the subgroup based on genotype. (B) Forest plot for associations between type 2 diabetes and *ApoE ε*2/ε3 genotype *vs*. ε3/ε3 genotype in the subgroup based on HWE.

**Supplementary Figure S4.**
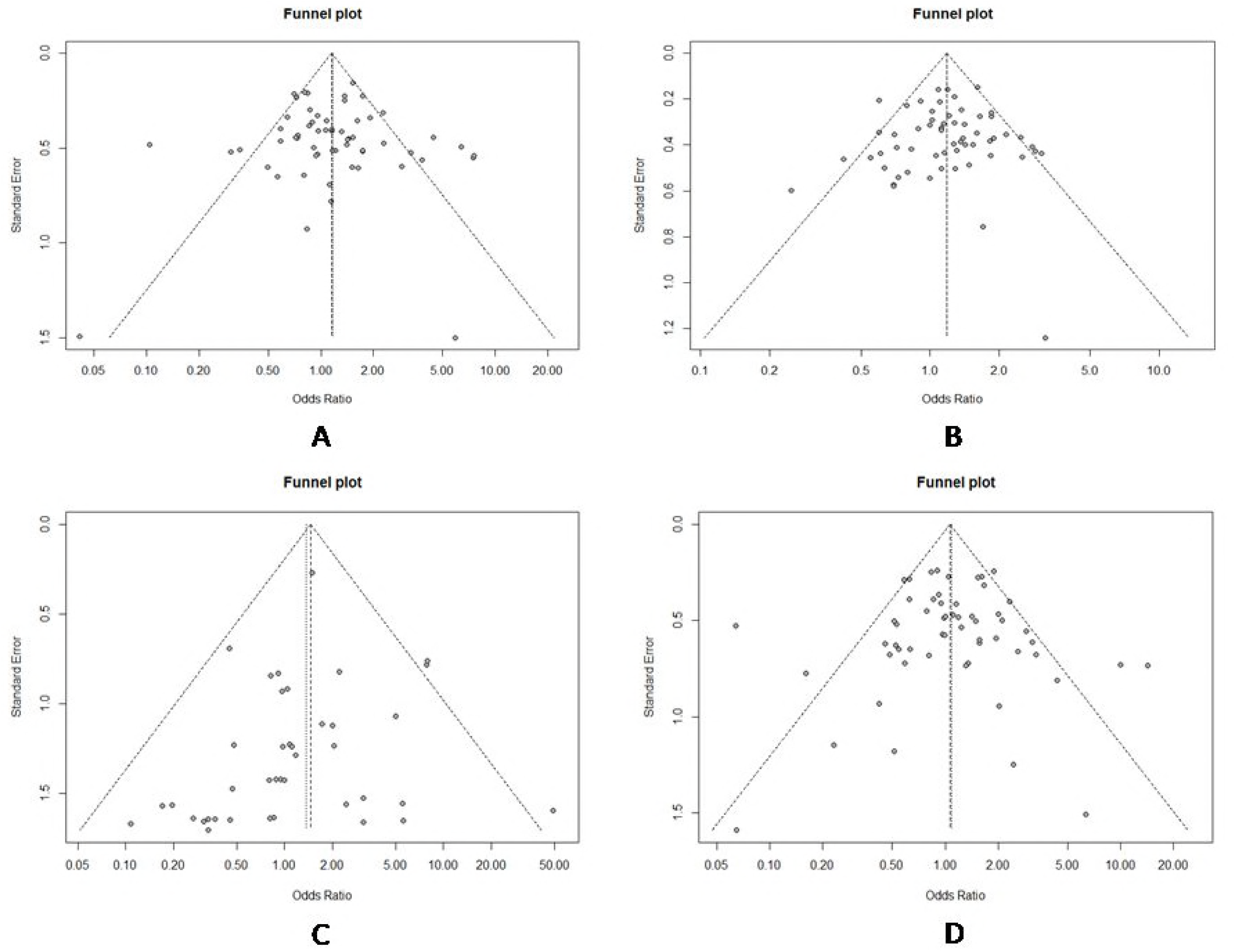
(A) Funnel plot of association between type 2 diabetes and *ApoE ε*2 allele *vs*. ε3 allele. (B) Funnel plot of association between type 2 diabetes and *ApoE ε*4 allele *vs*. ε3 allele. (C) Funnel plot of association between type 2 diabetes and *ApoE ε*2/ε2 genotype *vs*. and ε3/ε3 genotype. (D) Funnel plot of association between type 2 diabetes and *ApoE ε*2/ε3 genotype *vs*. and ε3/ε3 genotype.

**Supplementary Figure S5.**
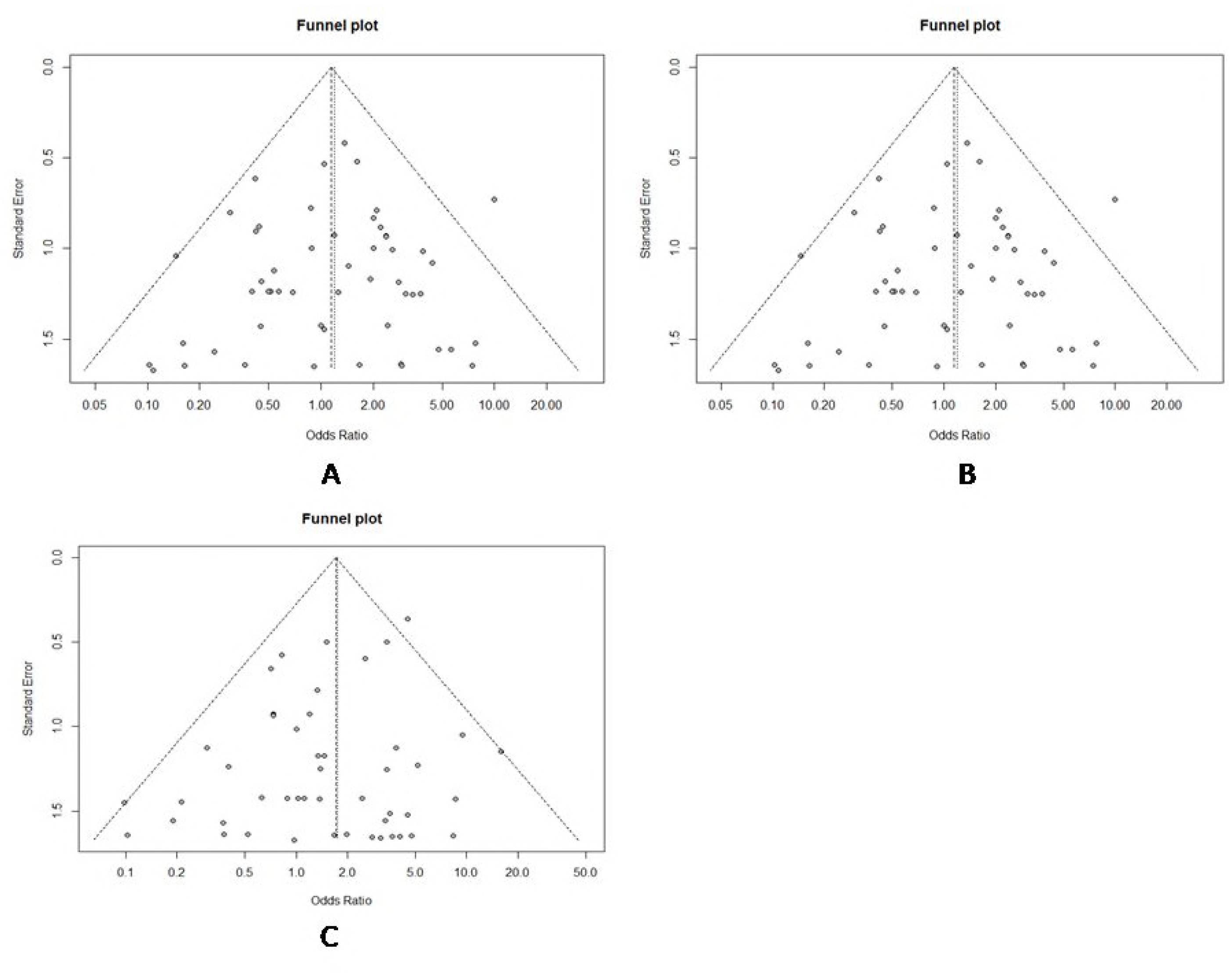
(A) Funnel plot of association between type 2 diabetes and *ApoE ε*2/ε4 genotype *vs*. and ε3/ε3 genotype. (B) Funnel plot of association between type 2 diabetes and *ApoE ε*3/ε4 genotype *vs*. and ε3/ε3 genotype.(C) Funnel plot of association between type 2 diabetes and *ApoE ε*4/ε4 genotype *vs*. and ε3/ε3 genotype.

